# TOPLESS promotes plant immunity by repressing auxin signaling and is targeted by the fungal effector Naked1

**DOI:** 10.1101/2021.05.04.442566

**Authors:** Fernando Navarrete, Michelle Gallei, Aleksandra E. Kornienko, Indira Saado, Khong-Sam Chia, Martin A. Darino, Mamoona Khan, Janos Bindics, Armin Djamei

## Abstract

In plants, the antagonism between growth and defense is hardwired by hormonal signaling. Perception of Pathogen Associated Molecular Patterns (PAMPs) from invading microorganisms inhibits auxin signaling and growth. Conversely, pathogens manipulate auxin signaling to promote disease, but how this hormone inhibits immunity is not fully understood. *Ustilago maydis* is a maize pathogen that induces auxin signaling in its host. We characterized an *U. maydis* effector protein, Naked1 (Nkd1), that is translocated into the host nucleus. Through its native EAR motif, Nkd1 binds to the transcriptional co-repressors TOPLESS/TOPLESS related (TPL/TPRs) and prevents the recruitment of transcriptional repressors involved in hormonal signaling, leading to de-repression of auxin and jasmonate signaling. Moderate up-regulation of auxin signaling inhibits an early defense response, PAMP-triggered ROS burst. Thus, our findings establish a clear mechanism for auxin-induced pathogen susceptibility. Manipulation of TPL by Nkd1 is evolutionary well balanced as engineered Nkd1 variants with increased expression or increased EAR-mediated TPL/TPR binding trigger typical salicylic acid mediated defense reactions, leading to pathogen resistance.

## Introduction

As sessile organisms, plants must integrate different endogenous and environmental signals to regulate growth and developmental programs. They have a limited set of resources that can be allocated either to growth or defense and pathogen attack usually leads to decreased growth and fitness. Thus, pathogen attack constitutes a fundamental factor negatively influencing plant growth and yield in agricultural systems (Bethany et al., 2014).

Plants sense the presence of invading microbes through a series of plasma membrane receptors called pattern recognition receptors (PRRs) (Dodds and Rathjen, 2010, Dangl and Jones, 2006). PRRs recognize highly conserved microbial molecules such as bacterial flagellin or fungal chitin. These molecules are collectively referred as pathogen/microbial associated molecular patterns (PAMPs/MAMPs). PRRs can also recognize endogenous molecules that are only produced after mechanical damage or pathogenic invasion, termed damage associated molecular patterns (DAMPs) (Boller and Felix, 2009). Independent of their origin, recognition of PAMPs /DAMPs triggers a series of stereotypic responses collectively termed pathogen/microbial triggered immunity (PTI/MTI). These include an increase in cytosolic Ca^2+^ levels, production of reactive oxygen species (ROS) by plasma membrane localized NADPH oxidases, activation of MAP kinases and transcriptional reprogramming, leading to growth inhibition (Dangl and Jones, 2006). PTI contributes to the inhibition of microbial colonization and therefore needs to be suppressed by pathogen secreted molecules, effectors, to establish the interaction. Effectors are thus vital for disease progression. Depending on the host genotype, effectors can be recognized by resistance (R) proteins, which frequently belong to the nucleotide-binding, leucine-rich-repeat (NLR) class. This recognition, termed effector triggered immunity (ETI), triggers a massive defense response, and is often characterized by a form of apoptotic cell death, the hypersensitive response (HR) (Dangl and Jones, 2006, Boller and Felix, 2009).

Since activation of defense responses is costly, it must be tightly regulated in order to optimize the tradeoffs between growth and defense (Belkhadir et al., 2014, Bethany et al., 2014). Hormonal crosstalk plays a major role in regulating this process and auxin, a growth-regulating hormone, has long been implicated in defense suppression. Despite being a well-studied phenomenon, the mechanisms of auxin-induced disease susceptibility are not fully understood. Many pathogens produce auxins to promote disease and exogenous application of this hormone has the same effect (Navarro et al., 2006, Ding et al., 2008, Suzuki et al., 2003, McClerklin et al., 2018). Auxin is thought to promote susceptibility in both salicylic acid-dependent (SA) and independent manners. Auxin treatment can prevent SA-dependent production of pathogenesis related proteins (PRs), which contribute to disease resistance (Aliab et al., 2018). This crosstalk is exploited by pathogenic bacteria. Besides synthesizing auxin, *P. syringae* uses the effector AvrRpt2 to target AUX/IAA repressor proteins, leading to increased auxin sensitivity and suppression of SA-dependent defenses (Chen et al., 2007, Cui et al., 2013). Auxin can also promote pathogen susceptibility independently of SA-mediated defenses, but the mechanism is unknown (Mutka et al., 2013). On the other hand, plants suppress auxin signaling following microbial recognition. Flagellin perception triggers the production of microRNA 393, which targets the auxin receptor TIR1, leading to repression of auxin signaling and increased resistance to *P. syringae* (Navarro et al., 2006).

Auxin is largely perceived in the nucleus. In the resting state, AUX/IAA repressors bind auxin response factors (ARFs) and repress transcription by recruiting the TOPLESS and TOPLESS-related (TPL/TPRs) co-repressors. TPL/TPRs are thought to act by recruiting histone deacetylases, therefore bringing the chromatin to a repressive state (Long et al., 2006, Szemenyei et al., 2008). In the presence of auxin, AUX/IAAs bind to TIR1/AFB receptors and are quickly ubiquitinated and degraded, subsequently releasing the repression of auxin-responsive genes (Mockaitis and Estelle, 2008).

TPL/TPRs mediate repression of other hormonal and developmental pathways in addition to auxin. Noticeably, jasmonate, brassinosteroids, and abscisic acid, which can also promote pathogen susceptibility (Pauwels et al., 2010, Causier et al., 2012, Oh et al., 2014, Lynch et al., 2017). Transcriptional repressors, like AUX/IAAs, use the short ethylene-responsive element binding factor-associated amphiphilic repression (EAR) motif to recruit TPL/TPRs. EAR motifs, with the amino acid sequence LxLxL, mediate the interaction with the N-terminal TOPLESS related domain (TRD) of TPL/TPRs (Ke et al., 2015, Martin-Arevalillo et al., 2017). Additionally, it has been shown that the interaction between the *Arabidopsis thaliana* TPR1 and the R protein SNC1 is necessary for activation of immune responses, although it seems to be independent of the presence of an EAR motif in SNC1 (Zhu et al., 2010).

*Ustilago maydis* is a fungal pathogen of maize. Infected plants form galls (cell masses derived from hypertrophy and hyperplasia) early after infection, which later become filled with fungal spores. Infected plants show an early, transient up-regulation and then suppression of genes involved in PTI, alongside the elevated expression of auxin, jasmonate, and gibberellin responsive genes (Doehlemann et al., 2008). Galls have elevated auxin levels (Turian and Hamilton, 1960). Although *U. maydis* can synthesize auxin, deletion of the necessary biosynthetic genes reduces auxin accumulation in galls but does not impair virulence (Reinecke et al., 2008). Thus, other factors, likely effector proteins, play a critical role in gall formation and virulence in this pathosystem. Effector encoding genes tend to form in clusters in the *U. maydis* genome, and their deletion can lead to significant reductions in virulence (Kämper et al., 2006, Schirawski et al., 2010, Navarrete et al., 2019).

Here we present the characterization of an effector from *U. maydis* found during a PTI inhibition screen. The protein, Naked1 (Nkd1), contains a functional EAR motif that mediates its interaction with members of the TOPLESS family of transcriptional co-repressors. Nkd1 is a virulence factor that acts in the maize nucleus. Nkd1 up-regulates jasmonate and auxin signaling, the latter of which leads to the suppression of PAMP-triggered ROS burst and increased pathogen susceptibility. We engineered variants of Nkd1 with altered affinity for members of the TOPLESS family and show that increased binding to TPL/TPRs leads to resistance reactions, likely due to elevated SA signaling. Using Nkd1 as a tool, our experiments support the notion that TPL/TPRs are conserved molecular nodes in plant growth-defense antagonism signaling.

## Results

### Nkd1 inhibits PAMP-triggered ROS burst in the plant nucleus

We identified Nkd1 (UMAG_02299) during a PTI inhibition screen of candidate effector proteins from *U. maydis*. We expressed the protein in *Nicotiana benthamiana* by Agrobacterium mediated transformation and tested its ability to suppress the PAMP-triggered ROS burst. Leaves expressing Nkd1_24-516_-mCherry-3xHA (lacking the predicted secretion signal) or mCherry (positive control) were challenged with chitin or flg22 and ROS production was monitored over time. Plants expressing Nkd1_24-516_-mCherry-3xHA showed a strongly reduced ROS burst compared with the mCherry control, independent of the PAMP used (Fig 1a). Next, we verified the localization of Nkd1 in *N. benthamiana* by confocal microscopy.

**Fig 1.**
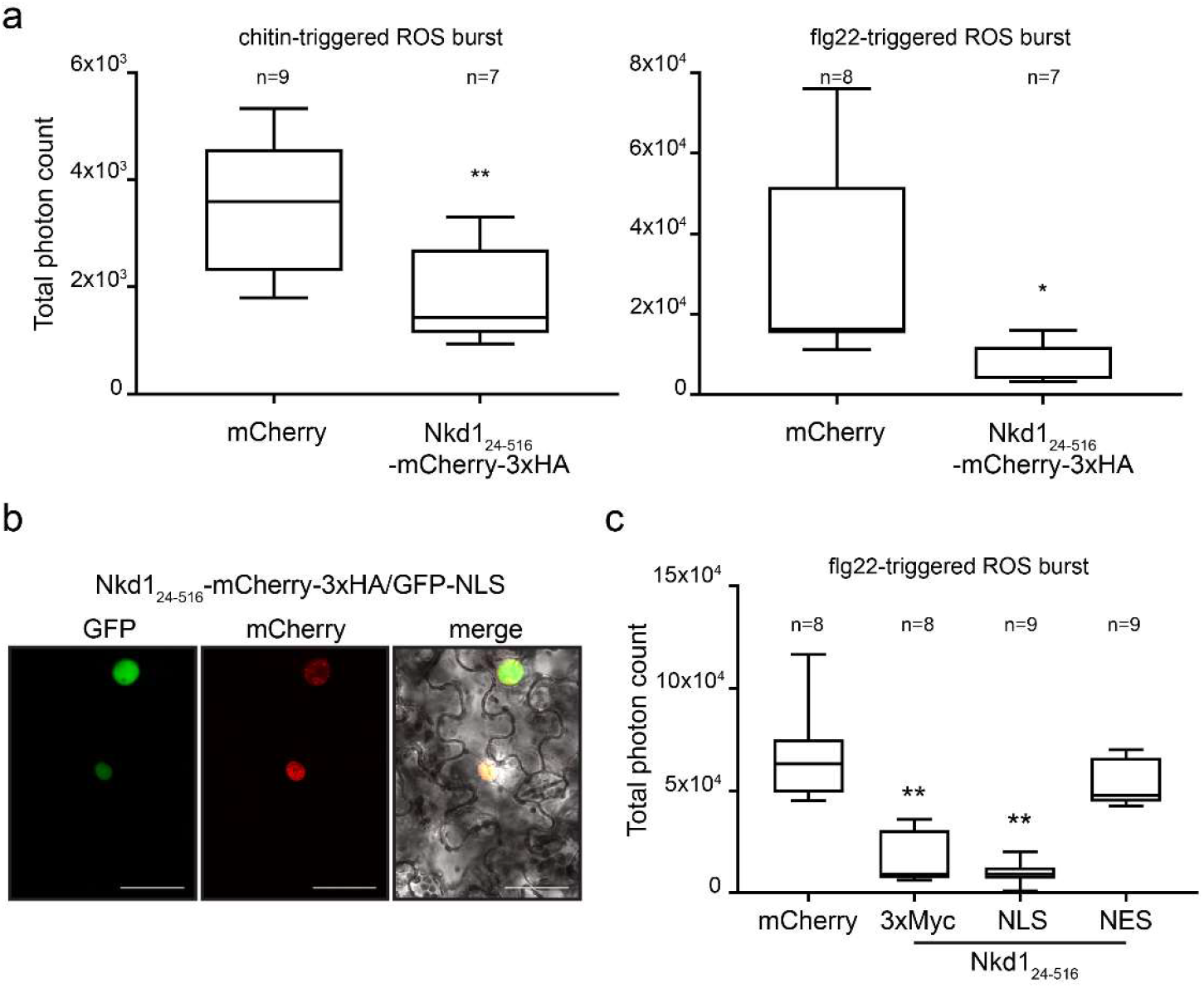
Nkd1 inhibits PAMP-triggered ROS burst in the plant nucleus. **a**. PAMP-triggered ROS burst in *N. benthamiana*. The effect of transient expression of Nkd1_24-516_-mCherry-3xHA on ROS burst triggered by chitin (left) or flg22 (right). Plants expressing mCherry were used as the positive control. Total photon counts indicating plant-derived apoplastic ROS were collected over 30 (chitin) or 40 minutes (flg22) and depicted as box plots. Data is a pool of three independent experiments (* p<0.05, ** p<0.01, t-test). **b**. Localization of NKd1 in *N. benthamiana*. Nkd1_24-516_-mCherry-3xHA was co-expressed with GFP-NLS in epidermal cells and localization was assessed by confocal microscopy. Nkd1_24-516_-mCherry-3xHA localizes to the nucleus. left panel: GFP fluorescence, middle panel: mCherry fluorescence, right panel: bright field-GFP-mCherry merge. Scale bar = 50 μm. **c**. Subcellular localization of Nkd1 affects ROS burst-suppressive activity. Nkd1_24-516_-NLS retains its ability to suppress flg22-triggered ROS burst whereas Nkd1_24-516_-NES does not. Data is a pool of three independent experiments. (* p<0.05, ** p<0.01, ANOVA, Tukeys). For all panels, n= number of plants used for each group.

Plants co-expressing Nkd1_24-516_-mCherry-3xHA and GFP-NLS (nuclear marker) showed that the effector protein localized exclusively to the nucleus (Fig 1b). We then verified whether the nuclear localization of Nkd1 is necessary for its immunity suppression activity using a mis-localization approach. We fused Nkd1 to either 3xMyc, an NLS (nuclear localization signal) or an NES (nuclear export signal), expressed the fusion proteins in *N. benthamiana* and measured PAMP-triggered ROS burst. 3xMyc was used as control since the Myc tag is not expected to alter protein localization. Plants expressing either Nkd1_24-516_-3xMyc or Nkd1_24-516_-NLS showed a very reduced ROS burst compared to the mCherry control whereas plants expressing Nkd1_24-516_-NES did not (Fig 1c). We confirmed the localization of the fusion proteins in *N. benthamiana* by confocal microscopy using Nkd1_24-516_-mCherry-3xHA, Nkd1_24-516_-mCherry-NLS and Nkd1_24-516_-mCherry-NES (Fig S1a). Effector fusions to mCherry-3xHA or mCherry-NLS showed reduced ROS burst compared to the control whereas the mCherry-NES fusion did not (Fig S1b). Therefore, our data indicates that Nkd1 suppresses immunity in the plant nucleus.

### Nkd1 promotes *U. maydis* virulence by targeting the maize nucleus

Since Nkd1 was originally found during a screen for putative effectors, we verified whether it shows the characteristics of effector proteins. *nkd1* is located on chromosome 5 in the previously identified effector cluster 5-21 (Schirawski et al., 2010) (Fig S2a). Analysis of *nkd1* expression by qPCR showed that it was induced 1-day post infection (dpi) of maize seedlings and peaked between 6 and 8 dpi (Fig S2b). Analysis of Nkd1 with SignalP 5.0 (Almagro Armenteros et al., 2019) predicted amino acids 1 through 23 to be a secretion signal. We therefore verified Nkd1 secretion by confocal microscopy during *U. maydis* infection in maize. We overexpressed the full length Nkd1fused to mCherry-3xHA (driven by the strong, biotrophy specific *pit2* promoter) and found that it localized to the edges and tips of the hyphae, indicating secretion (Fig 2a). We then plasmolyzed infected maize leaves, which expands the apoplastic space, and verified that Ndk1-mCherry-3xHA was freely diffusible in the apoplast (Fig S2c). In contrast, Nkd1_24-516_-mCherry-3xHA (lacking the predicted secretion signal) localized to the interior of the hyphae, indicating that secretion of Nkd1 depends on its predicted secretion signal (Fig 2a). Additionally, we verified the secretion of Nkd1 in axenic culture. We used the strain AB33, which filaments *in vitro*, mimicking some developmental changes undergone by *U. maydis* during host colonization (Brachmann et al., 2001). We could detect Nkd1_1-516_-3xHA in both, pellet and supernatant fractions, indicating that the protein is released form the hyphae, whereas the non-secrete actin control was only detected in the pellet fraction (Fig S2d).

**Fig 2.**
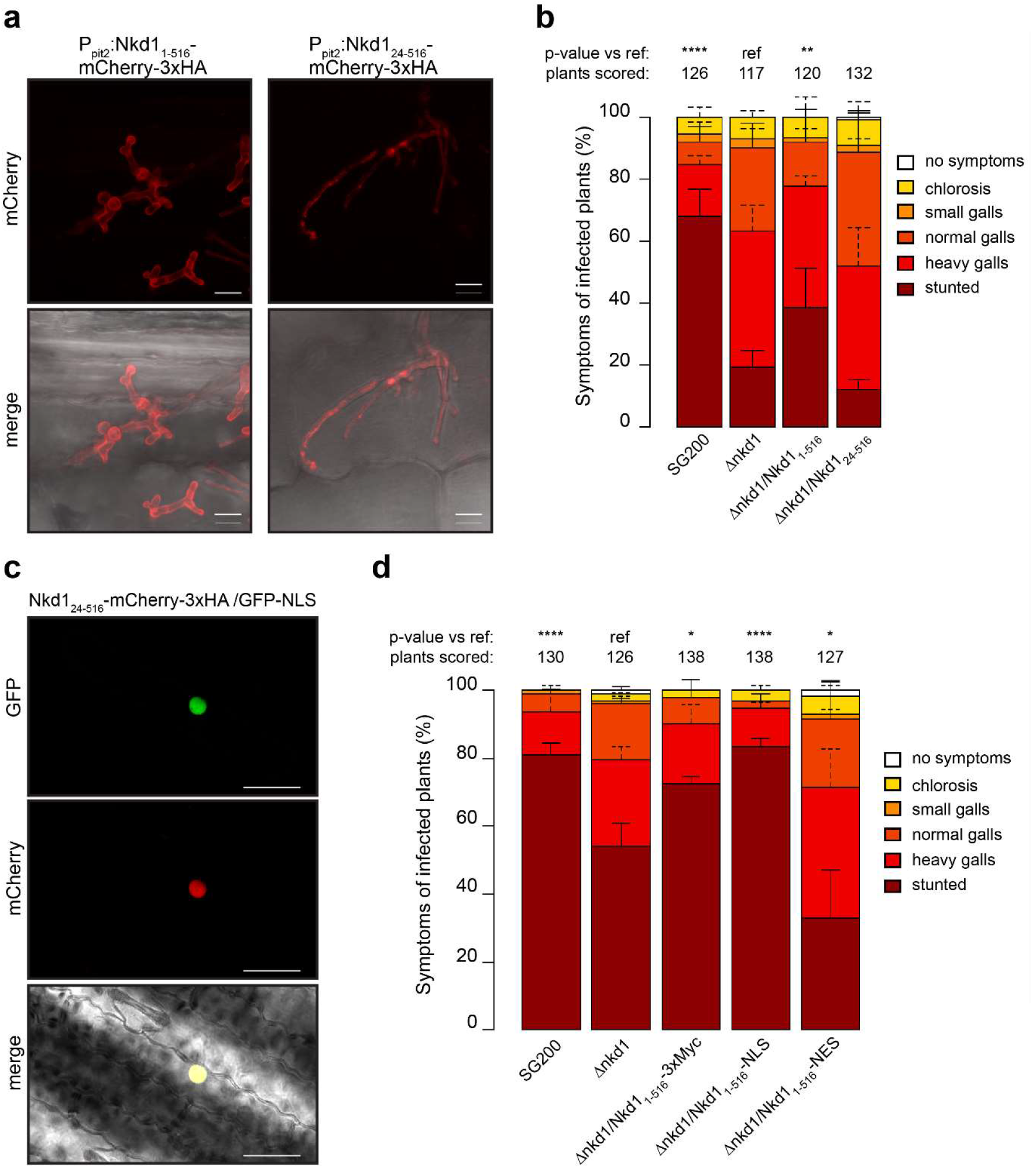
Nkd1 is a secreted protein that targets the host nucleus. **a**. Secretion of Nkd1 during maize infection. Left: *U. maydis* strain expressing P_pit2_:Nkd1_1-516_-mCherry-3xHA shows mCherry signal at the periphery of the hyphae and hyphal tips. Right: *U. maydis* strain expressing P_pit2_:Nkd1_24-516_-mCherry-3xHA (without its secretion signal) shows mCherry signal inside the hyphae. Pictures were taken 7 days post infection of maize seedlings. Upper panels: mCherry fluorescence, lower panels: bright field-mCherry merge. Scale bar = 10 μm. **b**. Disease symptom scoring of maize seedlings infected with *U. maydis* (7 dpi). SG200 (progenitor strain), ∆*nkd1* and two different complementation strains, either with (Nkd1_1-516_) or without (Nkd1_24-516_) the native secretion signal. **c**. Subcellular localization of Nkd1 in maize epidermal cells. Nkd1_24-516_-mCherry-3xHA was co-expressed with GFP-NLS in maize epidermal cells by biolistic bombardment. Nkd1_24-516_-mCherry-3xHA localizes to the nucleus. Upper panel: mCherry fluorescence, middle panel: GFP fluorescence, bottom panel: bright field-GFP-mCherry merge. Scale bar = 50 μm. **d**. Disease symptom scoring of maize seedlings infected with *U. maydis* (7 dpi). SG200 (progenitor strain), ∆*nkd1* and Nkd1 fused to tags that affect its localization, 3xMyc (neutral), NLS (nucleus), NES (cytoplasm). For b and d data represent mean ± SD from three independent experiments, n= total number of scored plants. Significant differences between strains were analyzed by the Fisher’s exact test with Benjamini-Hochberg correction for multiple comparisons (*p<0.05, ** p<0.01, *** p<0.001, **** p<0.0001).

We then generated deletion strains in the solopathogenic SG200 *U. maydis* background and infected maize seedlings to test whether *nkd1* contributes to virulence. Plants infected with the ∆*nkd1* strain showed milder symptoms compared with SG200 infected plants. Noticeably, stunting symptoms were much reduced in ∆*nkd1*-infected plants. (Fig 2b). Next, we tested whether secretion of Nkd1 contributes to virulence. We ectopically complemented the ∆*nkd1* mutant with constructs coding for the full-length protein (Nkd1_1-516_) or its derivative, lacking the predicted secretion signal (Nkd1_24-516_) and infected maize seedlings. Nkd1_1-516_ was able to partially complement the virulence defect of the ∆*nkd1* mutant whereas Nkd1_24-516_ was not, indicating that Nkd1 secretion is necessary for virulence (Fig 2b).

Since we showed earlier that Nkd1 inhibits defenses in the nucleus in *N. benthamiana*, we verified its sub-cellular localization in maize cells. We co-expressed 35S:Nkd1_24-516_-mCherry-3xHA with 35S:GFP-NLS in maize epidermal cells by biolistic bombardment. Confocal microscopy showed that Nkd1_24-516_-mCherry-3xHA co-localized with GFP-NLS in the maize nucleus and no mCherry signal could be detected in the cytoplasm (Fig 2c). We then tested whether localization to the maize nucleus is necessary for the virulence function of Nkd1. We employed a mis-localization approach identical to the one used for ROS burst suppression in *N. benthamiana*. We generated ∆*nkd1* complementation strains expressing full length Nkd1 fused to either 3xMyc, NLS, or NES. Expression of Nkd1_1-516_-3xMyc or Nkd1_1-516_-NLS complemented the virulence defect of the ∆*nkd1* mutant while Nkd1_1-516_-NES did not (Fig 2d), indicating that localization to the host nucleus is required for Nkd1 function. Fusing Nkd1_1-516_ to mCherry-3xHA was not able to rescue the virulence defect of the ∆*nkd1* mutant (Fig S2e), even though Nkd1_24-516_-mCherry-3xHA was active in plant cells (Fig S1). Since fusion to large, structurally stable proteins like GFP or mCherry has been reported to inhibit effector translocation, but not secretion (Tanaka et al., 2015), we interpret this result as further indication that Nkd1 is translocated into the host cells.

Taking together, our results from *N. benthamiana* and maize infection experiments indicate that Nkd1 is secreted from *U. maydis* and translocated into the host nucleus, where it promotes virulence. Further, our data also imply that the target(s) of Nkd1 must be conserved between monocots and dicots.

### Nkd1 binds to TPL/TPRs to suppress the PAMP-triggered ROS burst

In order to gain further insight into the function of Nkd1, we performed homology searches using BLAST. We found homologs of UmNkd1 only in the closely related smut fungi *Pseudozyma hubeiensis, Sporisorium reilianum, S. scitamineum, S. graminicola*, and *Melanopsichium pennsylvanicum*. A close examination of the C-terminal sequence of these proteins revealed the presence of an LxLxL-type EAR motif that was conserved across most of the orthologs. In *S. scitamineum*, the second L residue is substituted by the structurally related I, and in *M. pennsylvanicum* (which shows the least sequence homology to UmNkd1 overall) the motif is missing completely (Fig 3a). We therefore evaluated whether the presence of an EAR motif is important for the function of the Nkd1 orthologs. We expressed UmNkd1, SrNkd1, SsNkd1, and MpNkd1 without their predicted secretion signals in *N. benthamiana* and tested their ability to suppress immunity as before. All orthologs were expressed as full-length proteins and localized to the nucleus (Fig S3a, b) but only MpNkd1, which lacks the EAR motif, failed to suppress the PAMP-triggered ROS burst (Fig 3b). This result indicates that the EAR motif could be required for this process.

**Fig 3.**
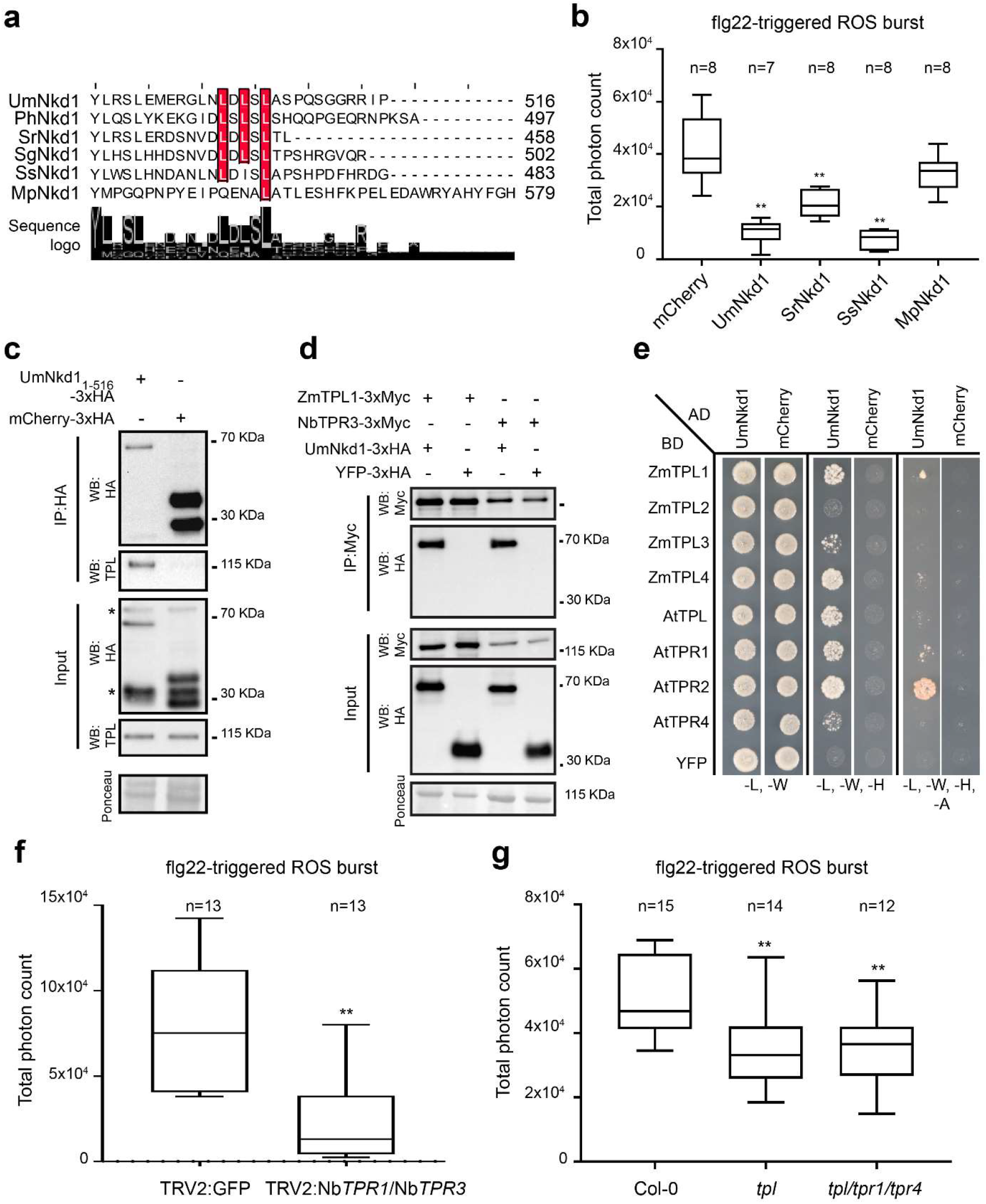
Nkd1 interacts with members of the TOPLESS family. **a**. Protein alignment of the C-terminal portion of Nkd1 and its orthologs (Um: *U. maydis*, Ph: *Pseudozyma hubensis*, Sr: *Sporisorium rellianum*, Sg: *Sporisorium graminicola*, Ss: *Sporisorium scitamineum*, Mp: *Melanopsichium pennsylvanicum*). Conserved leucine residues corresponding to the EAR motif are highlighted in red. Numbers on the right indicate protein length in aa. **b**. PAMP-triggered ROS burst suppression in *N. benthamiana* expressing different Nkd1 orthologs. Only MpNkd1, which lacks the EAR motif, is unable to suppress the PAMP-triggered ROS burst. Plants expressing mCherry were used as the positive control. Total photon counts over 40 min are shown as box plots. Data is a pool of three independent experiments (** p<0.01, ANOVA, Tukeys). **c**. UmNkd1 interacts with TPL during maize infection. Maize plants were infected with *U. maydis* strain carrying P_Pit2_:Nkd1_1-516_-3xHA or P_Pit2_:Nkd1_1-23_-mCherry-3xHA. Total proteins were immunoprecipitated with α-HA magnetic beads (IP: HA) and blotted with specific antibodies. ZmTPLs co-immunoprecipitate with Nkd1_1-516_-3xHA but not with mCherry-3xHA. *: unspecific band. **d**. UmNkd1 interacts with members of the TPL family in *N. benthamiana*. Total proteins were immunoprecipitated with α-Myc magnetic beads (IP: Myc) and blotted with specific antibodies. UmNkd1_24-516_-3xHA but not YFP-3xHA, co-immunoprecipitates with ZmTPL1-3xMyc and NbTPR3-3xMyc. **e**. Y2H assay showing interaction of UmNkd1_24-516_ with TPL homologs. Yeast strains were grown on SD medium lacking indicated amino acids/nucleotides. Growth in media lacking leu (L) and trp (W) is used as transformation control. Growth in intermediate stringency media lacking leu (L), trp (W), and his (H) or high stringency media lacking leu (L), trp (W), his (H), and ade (A) indicates protein interaction. **f**. flg22-triggered ROS burst in *N. benthamiana* plants in which *TPR1* and *TPR3* were silenced by VIGS, TRV2:GFP was used as an off-target control. **g**. flg22-triggered ROS burst in *A. thaliana* Col-0 and *tpl* mutants. For f and g, total photon counts over 40 min are shown as box plots. Data is a pool of three independent experiments (** p<0.01, Tukey’s ANOVA). For all panels, n= number of plants used for each group.

Since the EAR motif is known to mediate protein interactions with TPL/TPRs in plants (Causier et al., 2012), these results prompted us to investigate whether UmNkd1 interacts with members of this family. We constructed *U. maydis* strains expressing UmNkd1-3xHA or mCherry-3xHA driven by the Um*pit2* promoter, which confers strong expression during biotrophic development (Doehlemann et al., 2011, Lanver et al., 2018) and used them to infect maize seedlings. We extracted total proteins from infected tissues and immunoprecipitated HA-tagged proteins with α-HA magnetic beads. By western blot with α-TPL antibodies we showed that TPL proteins co-immunoprecipitated (Co-IP) with UmNkd1-3xHA but not with mCherry-3xHA (Fig 3c), which indicates interaction between TPLs and the effector protein. To confirm this interaction, we co-expressed ZmTPL1-3xMyc and NbTPR3-3xMyc with UmNkd1_24-516_-3xHA or YFP-3xHA in *N. benthamiana* and immunoprecipitated TPL/TPRs with α-Myc magnetic beads. UmNkd1_24-516_-3xHA, but not YFP-3xHA, co-immunoprecipitated with ZmTPL1-3xMyc and NbTPR3-3xMyc, confirming our previous results (Fig 3d). Further, to test whether the interaction between UmNkd1 and TPL/TPRs is direct, we performed a yeast two-hybrid assays (Y2H). TPL/TPR homologs from maize and *A. thaliana* were fused with the GAL4BD and used to test their interaction with UmNkd1_24-516_ or mCherry fused with the GAL4AD. UmNkd1 interacted with all homologs tested with the exception of ZmTPL2 and it seems to show preference for ZmTPL1/ZmTPL4 and for AtTPR2/AtTPL/AtTPL1 as evidenced by the differential growth on the different selective media (Fig 3e). The implications of this apparent preference for certain homologs might indicate sub-specialization of the different TPL/TPRs but it awaits further studies.

The interaction between UmNkd1 and TPL/TPR suggested that these plant proteins are involved in PTI, a function not previously attributed to them. Therefore, we tested this hypothesis by silencing *TPRs* in *N. benthamiana* by virus-induced gene silencing and assessing *tpl/tpr* mutants in *A. thaliana. NbTPR1*/*TPR3*-silenced plants and single (*tpl*) or triple (*tpl/tpr1/tpr4*) *A. thaliana* knockout plants showed a reduced PAMP-triggered ROS burst compared to their controls (Fig 3f, g). Therefore, taken together, our data shows that TPL/TPR co-repressors regulate PAMP-triggered ROS burst and that Nkd1 proteins target TPL/TPRs to suppress this process.

### The Nkd1 EAR motif mediates its binding to TPL/TPRs and de-repression of hormonal signaling

Since we established that the EAR motif of the Nkd1 family is necessary for immune suppression, we mutated this motif in UmNkd1 and evaluated its effect on the binding to TPL/TPRs. Three mutants were made, Nkd1EARm (L501A, alanine substitution in the first position of the conserved EAR motif), Nkd1∆EAR (deletion of the last 16 amino acids, including the EAR motif) and Ndk1SRDX (Nkd1∆EAR fused to SRDX, an optimized EAR motif derived from the SUPERMAN transcription factor) (Hiratsu et al., 2003)(Fig 4a). By Y2H assays we showed that Nkd1EARm had reduced binding to TPL/TPRs from maize and *A. thaliana*. Only minor growth was visible in intermediate stringency media (-L, -W, -H) compared to the WT control. Further, Nkd1∆EAR did not bind to any of the TPL/TPRs, as indicated by the lack of growth on intermediate stringency media. In contrast, Ndk1SRDX showed increased binding to TPL/TPRs compared to the WT control, indicated by strong growth in both intermediate and high stringency media (-L, -W, -H, -A). In addition, this mutant was the only one that showed interaction with ZmTPL2 (Fig 4b). We then confirmed these results by CoIP in *N. benthamiana*. We co-expressed ZmTPL1-3xMyc with either Nkd1-3xHA, Nkd1EARm-3xHA, Nkd1∆EAR-3xHA, Ndk1SRDX-3xHA, or YFP-3xHA and immunoprecipitated total proteins with α-Myc magnetic beads. Nkd1EARm-3xHA and Nkd1∆EAR-3xHA showed strongly reduced binding to ZmTPL1-3xMyc compared to the WT control, as indicated by the strongly reduced or absent signal in the α-HA blot of the IP fraction (Fig 4c). On the other hand, Ndk1SRDX-3xHA co-precipitated with ZmTPL1-3xMyc and showed stronger binding compared to the WT control, as indicated by the increased signal in the α-HA blot of the IP fraction (Fig 4c). Our results thus indicate that Nkd1 interacts with TPL/TPRs through its EAR motif and that the specific sequence of this motif determines the strength of the interaction.

**Fig 4.**
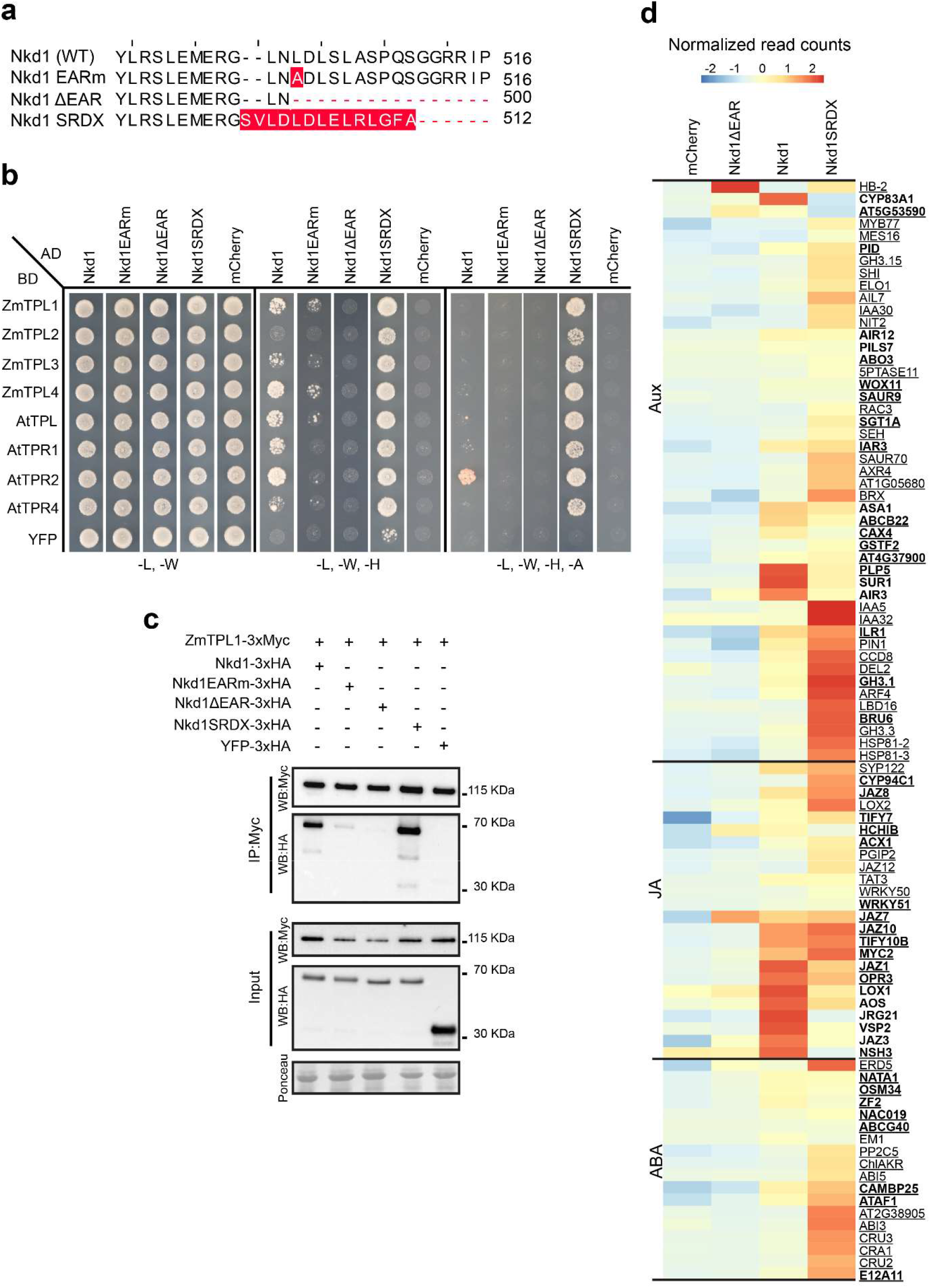
The EAR motif of Nkd1 mediates binding to TPL/TPRs and de-repression of hormone signaling. **a**. Protein alignment of the C-terminal portion of UmNkd1 and its EAR mutants. For each protein, the mutated residues are highlighted in red. Numbers on the right indicate protein length in aa. **b**. Y2H assay showing interaction of Nkd1 and its EAR mutants with TPL homologs. Yeast strains were grown on SD medium lacking indicated amino acids/nucleotides. Growth in media lacking leu (L) and trp (W) is used as transformation control. Growth in intermediate stringency media lacking leu (L), trp (W), and his (H) or high stringency media lacking leu (L), trp (W), his (H), and ade (A) indicates protein interaction. **c**. Interaction of Nkd1 and its EAR mutants with ZmTPL1 *in planta*. Total proteins were immunoprecipitated from *N. benthamiana* with α-Myc magnetic beads (IP: Myc) and blotted with specific antibodies. Nkd1-EARm and Nkd1 ΔEAR show reduced affinity to ZmTPL1 whereas Nkd1SRDX shows increased affinity to ZmTPL1. **d**. Heat map showing the expression levels of hormone responsive genes in *A. thaliana* seedlings 5h post induction of transgene expression. Aux: auxin, JA: jasmonic acid, ABA: abscisic acid, responsive/homeostatic genes. Genes depicted here are differentially expressed in at least one of the lines compared to the mCherry expressing plants (control). Bold: significantly miss-regulated in Nkd1. Underlined: significantly miss-regulated in Nkd1SRDX. Data is the average of 3 independent experiments. Differential gene expression was analyzed with DESeq2, log2Fold change >0.6, p <0.01 (Benjamini-Hochberg correction).

Taking advantage of the fact that UmNkd1 binds to TPL/TPRs from different species, we generated transgenic *A. thaliana* plants expressing the effector protein under the control of an estradiol inducible promoter. *A. thaliana* seedlings expressing either Nkd1-3xMyc, Nkd1∆EAR-3xMyc, Nkd1SRDX-3xMyc, or mCherry-3xMyc (reference control) were treated with estradiol for 5 and 24 h and gene expression was monitored by RNAseq. We observed transcriptomic changes that varied according to the strength of the interaction between the different Nkd1 variants and TPL/TPRs. The number of differentially expressed genes (DEGs) was higher in plants expressing Nkd1SRDX-3xMyc than in plants expressing Nkd1-3xMyc and Nkd1∆EAR-3xMyc (Fig S4, Fig 4d, Supplementary Table 1). For instance, 5 h after estradiol treatment we detected 2156, 910 and 21 upregulated genes in Nkd1SRDX-3xMyc, Nkd1-3xMyc and Nkd1∆EAR-3xMyc expressing plants, respectively (Fig S4a). Since the latter plants showed so few DEGs this indicates that most, if not all physiological changes induced by the expression of Nkd1 in *A. thaliana* derive from EAR-mediated interactions with plant proteins. GO term analysis of DEGs identified 5 h after estradiol treatment showed that most upregulated genes belonged to the classes “defense response”, “response to other organism”, and “immune response”. The enrichment for these categories was several orders of magnitude higher in plants expressing Nkd1SRDX-3xMyc than in plants expressing Nkd1-3xMyc (Fig S4b). These categories contain many of the genes involved in PTI responses such as *RLKs, MAPKs*, and the plasma membrane NADPH oxidase *RBOHD*, which is responsible for the PAMP-triggered ROS burst, and therefore indicate that immunity suppression by Nkd1 is most likely post-transcriptional.

We then examined other GO categories (with fewer DEGs) that could explain susceptibility towards biotrophs. These included, “response to endogenous stimulus”, “cellular response to jasmonic acid stimulus”, “cellular response to hypoxia”, and “response to hormone”. These pathways contain genes whose transcriptional regulation is mediated by TPL/TPRs and rely on transcriptional repressors with EAR motifs of the same type found in Nkd1 (Szemenyei et al., 2008, Pauwels et al., 2010, Causier et al., 2012, Lynch et al., 2017). Lines expressing Nkd1-3xMyc and Nkd1SRDX-3xMyc showed upregulation of genes involved in the signaling and metabolism of auxin and jasmonic acid. We observed upregulation of typical early auxin-responsive genes such as *GH3*.1, *BRU6* and *SAUR9* in plants expressing either Nkd1-3xMyc or Nkd1SRDX-3xMyc, the latter also showed up-regulation of *AUX/IAA* repressors and the *PIN1* auxin transporter (Fig 4d, Supplementary Table 1). Plants expressing either Nkd1-3xMyc or Nkd1SRDX-3xMyc also showed upregulation of typical early jasmonic-responsive genes, such as biosynthetic genes and *JAZ* transcriptional repressors (Fig 4d, Supplementary Table 1). In addition, plants expressing Nkd1SRDX-3xMyc showed stronger upregulation of abscisic acid responsive genes compared to plants expressing Nkd1 (Fig 4d, Supplementary Table 1).

Overall, our results indicate that the specific sequence of the EAR motif of Nkd1 determines the strength of its interaction with TPL/TPRs and the degree to which the latter can repress gene expression.

### TPL/TPRs mediated repression of auxin signaling is required for PAMP-triggered ROS burst

The fact that EAR-mediated interaction between Nkd1 and TPL/TPRs leads to the up-regulation of TPL/TPR repressed genes led us to hypothesize that Nkd1 can affect the recruitment of TPL/TPRs co-repressors to transcriptional repressors that contain EAR motifs. We tested this hypothesis by assessing the ability of AUX/IAA proteins to bind to TPL/TPRs in the presence of Nkd1. We co-expressed ZmTPL1-3xMyc and ZmIAA5-GFP in the presence of either Nkd1-3xHA, Nkd1∆EAR-3xHA, Nkd1SRDX-3xHA, or mCherry-3xHA in *N. benthamiana* and immunoprecipitated ZmTPL1 with α-Myc magnetic beads. ZmIAA5-GFP was able to co-precipitate with ZmTPL1-3xMyc in the presence of mCherry-3xHA or Nkd1∆EAR-3xHA but not in the presence of Nkd1-3xHA or Nkd1SRDX-3xHA (Fig 5a), indicating that EAR motif-mediated binding of Nkd1 to TPL/TPRs prevents its recruitment to AUX/IAAs and likely mediates the up regulation of hormone responsive genes. Consistent with this result, we co-expressed 35S:Nkd1_24-516_-mCherry-3xHA or 35S:mCherry with the DR5:YFP auxin reporter in maize epidermal cells by biolistic bombardment. Nkd1-mCherry-3xHA was able to up-regulate the expression of the DR5:YFP reporter compared with the mCherry control. (Fig 5b). Furthermore, Nkd1-3xMyc and Nkd1SRDX-3xMyc but not Nkd1EARm-3xMyc or Nkd1∆EAR-3xMyc were able to up-regulate the expression of the DR5:GFP reporter in *A. thaliana* and *N. benthamiana* (Fig 5e, S5c). Additionally, UmNkd1, SrNkd1, and SsNkd1, but not MpNkd1 (which lacks an EAR motif), were able to upregulate expression of the DR5 reporter upon transient expression in *N. benthamiana* (Fig S5d).

**Fig 5.**
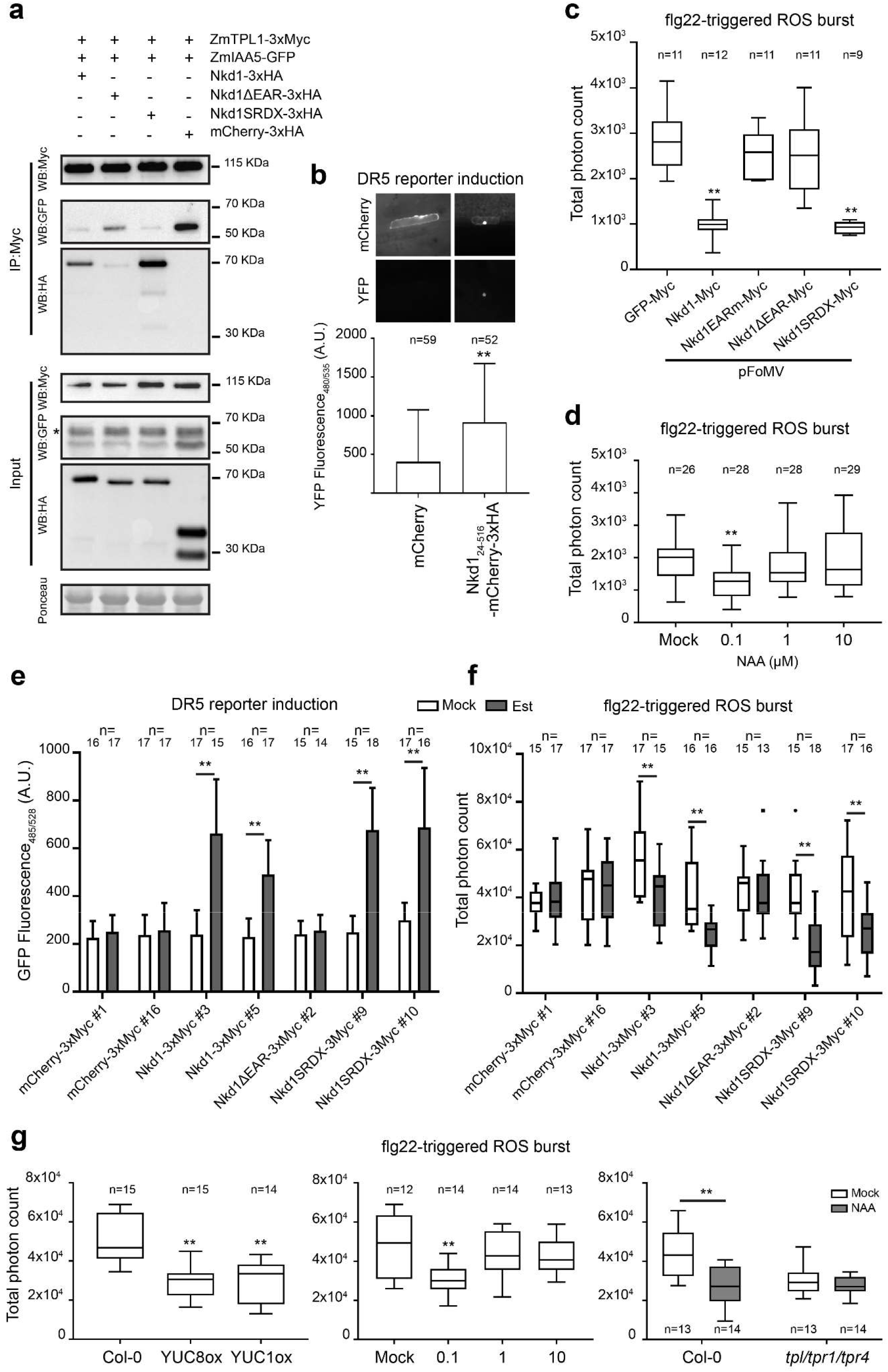
TPL/TPR mediated repression of auxin signaling is required for PAMP-triggered ROS burst. **a**. Proteins were immunoprecipitated from *N. benthamiana* with α-Myc magnetic beads (IP: Myc) and blotted with specific antibodies. ZmIAA5 co-precipitates with ZmTPL1 in the presence of Nkd1ΔEAR or mCherry but not in the presence of Nkd1 or Nkd1SRDX. * un-specific band. **b**. Nkd1_24-516_-mCherry-3xHA or mCherry were co-expressed with DR5:YFP in maize epidermal cells. Top: representative pictures of cells expressing either mCherry or Nkd1_24-516_-mCherry-3xHA. Bottom: reporter activity quantified by fluorescence intensity (528 nm) of transformed cells. Data is a pool of 3 independent experiments, mean ± SD (** p<0.01, t-test). **c**. Nkd1 variants were expressed transiently in maize with the pFoMV viral vector. Nkd1 and Nkd1SRDX but not Nkd1EARm or Nkd1∆EAR inhibit flg22-triggered ROS burst 9 dpi. Data (total photon counts over 20 min) is a pool of 3 independent experiments (** p<0.01, Tukey’s ANOVA). **d**. Maize plants treated with 0.1 µM NAA show reduced ROS burst compared to the mock treatment. Data is a pool of four independent experiments (** p<0.01 Tukeys’s ANOVA). **e**. Nkd1 and Nkd1SRDX, but not Nkd1∆EAR, up-regulate the expression of the DR5:GFP reporter in *A. thaliana*. Data is a pool of 3 independent experiments, mean ± SD (** p<0.01, Tukey’s ANOVA). **f**. Nkd1 and Nkd1SRDX but not Nkd1∆EAR, inhibit PAMP triggered ROS burst in *A. thaliana*. Data (total photon counts over 40 min) is a pool of 3 independent experiments (** p<0.01, Bonferroni’s Two way-ANOVA). Measurements were done 24h after β-estradiol treatment. **g**. Auxin inhibits flg22-trigered ROS burst in *A. thaliana*. Left: *A. thaliana* plants overexpressing YUCCA proteins have reduced flg22-triggered ROS burst compared to the Col-0. Center: plants treated with 0.1 µM NAA show reduced ROS burst compared to the mock treatment. Right: treatment of plants with 0.1 µM NAA inhibits the ROS burst in WT but not in the *tpl/tpr1/tpr4* triple mutant. Data (total photon counts over 40 min) is a pool of 3 independent experiments (** p<0.01, left/center: Tukey’s ANOVA, right: Bonferroni’s two way-ANOVA). For all panels, n= number of samples used for each group.

We also evaluated the effect of the EAR mutations on the ability of Nkd1 to suppress the PAMP-triggered ROS burst. We transiently expressed different Nkd1 variants in maize by using vectors derived from the Foxtail mosaic virus (pFoMV) (Bouton et al., 2018). Parallel to our observations with the DR5 reporter, Nkd1 and Nkd1SRDX but not Nkd1EARm or Nkd1∆EAR, were able to suppress PAMP-triggered ROS burst in maize plants compared to the GFP control (Fig 5c). This was despite the fact that constructs encoding different Nkd1 variants yielded uneven viral loads and subsequently different protein amounts in maize (Fig S5a, S5b). We confirmed this result by expressing the same effector variants in *A. thaliana* and *N. benthamiana* (Fig 5f, S5e). Our results thus indicate that either Nkd1 affects auxin signaling and PAMP-triggered ROS simultaneously or that up-regulation of auxin signaling could suppress PAMP-triggered ROS burst. Moreover, the phenomenon seems to be conserved across monocot and dicot plants.

To test this hypothesis, we assessed PTI in *YUC8* and *YUC1* overexpressing lines in *A. thaliana* plants (which have high endogenous levels of auxin)(Zhao et al., 2001, Hentrich et al., 2013). Both lines showed a reduced PAMP-triggered ROS burst compared to the wild-type Col-0 background (Fig 5g). Furthermore, treatment of maize or *A. thaliana* plants with 100 nM of the auxin analog NAA, but not higher concentrations, led to a reduction in PAMP-triggered ROS burst (Fig 5d, 5g). Finally, we evaluated the effect of *tpl* mutations on the auxin-mediated ROS burst inhibition. NAA treatment led to a reduced PAMP-triggered ROS burst in Col-0 but not in the *tpl/tpr1/tpr4* knockout, indicating that TPL/TPRs are necessary for auxin-mediated suppression of the PAMP-triggered ROS burst.

Overall, our results indicate that binding of Nkd1 to TPL/TPRs prevents recruitment of AUX/IAAs, and possibly other transcriptional repressors, leading to an up-regulation of auxin signaling and a subsequent suppression of the PAMP-triggered ROS burst.

### Increased Nkd1 binding to TPL/TPRs triggers resistance reactions

In order to evaluate how the binding of Nkd1 to TPL/TPRs affects the virulence of *U. maydis*, we ectopically complemented the ∆*nkd1* mutant with constructs coding for the WT protein or its EAR mutants (Nkd1EARm, Nkd1ΔEAR, and Nkd1SRDX). Remarkably, Nkd1EARm and Nkd1∆EAR complemented the virulence defect of the ∆*nkd1* mutant, similar to the levels observed with the WT *nkd1*. Even more unexpectedly, complementing ∆*nkd1* with Nkd1SRDX led to an even more pronounced virulence defect than was observed in the knockout (Fig 6a). A similar phenotype was observed when dramatically increasing the expression of *nkd1* during infection in maize. Complementation of the ∆*nkd1* with P_Cmu1_:nkd1 (the *cmu1* promoter has a very high expression during infection, Djamei et al., 2011) also lead to a more pronounced virulence defect than what was observed in the knockout (Fig 6b). We then used propidium iodide, which stains cells with compromised plasma membrane and cell walls to assess host cell viability during infection with the different strains (Doehlemann et al., 2009, Jones et al., 2016). Maize plants infected with either ∆*nkd1/* P_Nkd1_:nkd1-SRDX or ∆*nkd1/* P_Cmu1_:nkd1 showed cell death in cells surrounding invading hyphae, whereas this was not observed in plants infected with SG200, ∆*nkd1* or ∆*nkd1*/nkd1 (Fig S6), indicating that the pronounced virulence defect of the former two strains is due to increased cell death/immune reactions.

**Figure 6.**
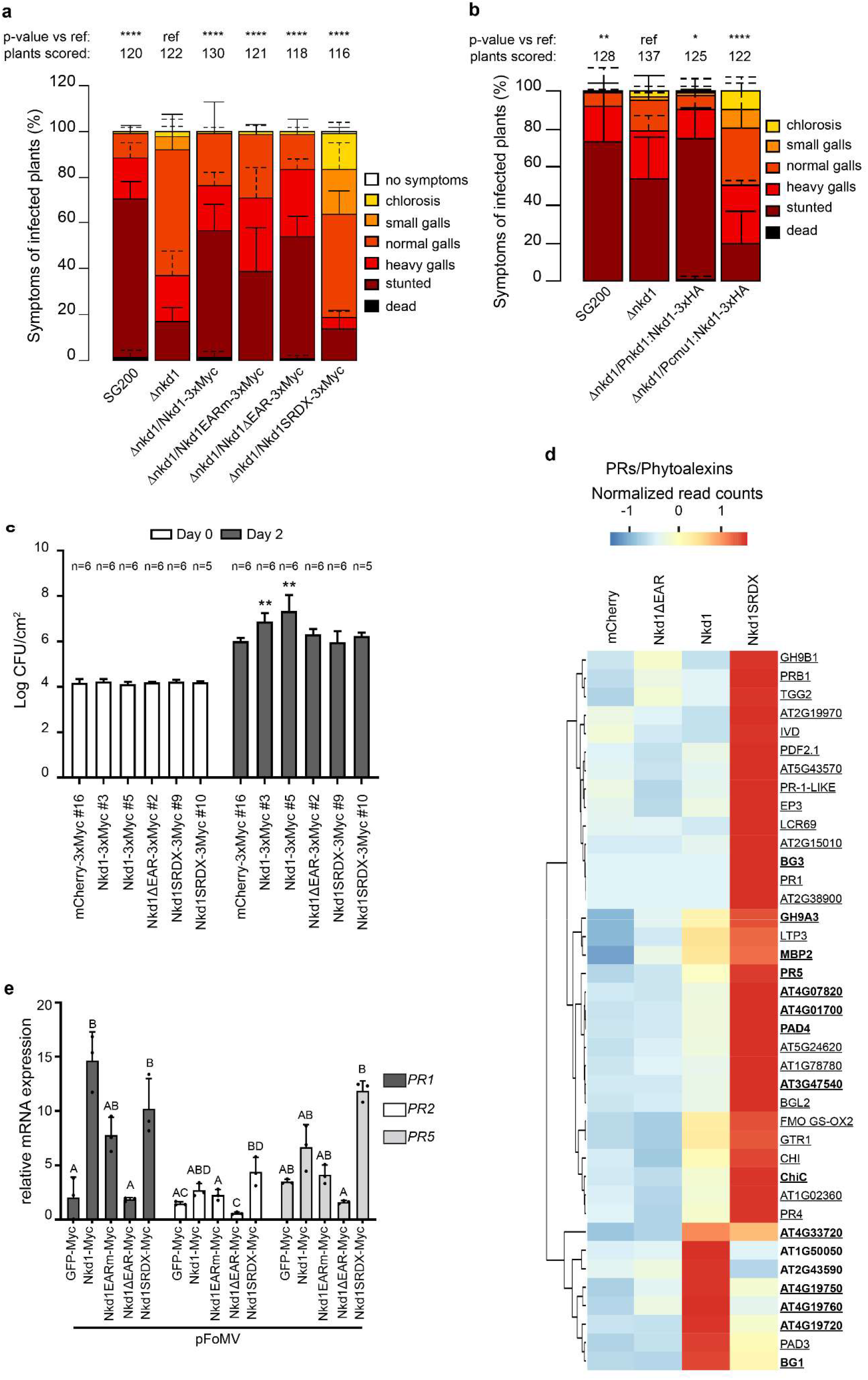
The EAR motif of Nkd1 affect host susceptibility to pathogens. **a**. Disease symptom scoring of maize seedlings infected with *U. maydis*. SG200 (progenitor strain), ∆*nkd1* and different complementation strains carrying mutations in the EAR motif (7dpi). Nkd1 (WT), Nkd1EARm, and Nkd1ΔEAR are able to complement the virulence defect of *U. maydis* Δnkd1 but Nkd1SRDX is not. Data represent mean ± SD from three independent experiments, n= total number of scored plants. Significant differences between strains were analyzed by the Fisher’s exact test with Benjamini-Hochberg correction for multiple comparisons (*p<0.05, ** p<0.01, *** p<0.001, **** p<0.0001). **b**. Bacterial growth in *A. thaliana* expressing different Nkd1 constructs. Plants expressing mCherry were used as the control. Nkd1 (WT) promotes susceptibility of *A. thaliana* to *P. syringae* but Nkd1ΔEAR and Nkd1SRDX do not. Data is one representative experiment from a total of 3, n= number of plants used for each group. Significant differences between lines were analyzed by Tukey’s ANOVA, (** p<0.01). **c**. Heat map showing the expression levels of PRs and phytoalexin biosynthetic and regulatory genes in *A. thaliana* seedlings 24h post induction of trans gene expression (Euclidian clustering). Genes depicted are differentially expressed in at least one of the lines compared to the mCherry expressing plants (control). Bold: significantly upregulated in Nkd1. Underlined: significantly upregulated in Nkd1SRDX. Data is the average of 3 independent experiments. Genes significantly upregulated in both Nkd1 and Nkd1SRDX lines are highlighted in bold. Data is the average of 3 independent experiments. Differential gene exp was analyzed with DESeq2, log2Fold change >0.6, p <0.01 (Benjamini-Hochberg correction). **d**. *PR* gene expression in maize plants expressing the respective viral constructs was monitored by qRT-PCR and normalized to Zm*CDK* 9dpi. Data is mean ± SD, n=3 (different letters indicate statistical difference, p<0.01, by Tukey’s ANOVA).

To gain insights into how plants react to Nkd1 beyond the suppression of PAMP triggered ROS burst, we infected *A. thaliana* lines expressing Nkd1 or its EAR mutants with *Pseudomonas syringae* pv. tomato DC3000 (Pst) and evaluated bacterial colonization.

Expression of WT Nkd1 led to increased susceptibility to Pst. These lines showed 6 to 30-fold increase in bacterial populations compared to plants expressing mCherry (reference control). In contrast, bacterial counts in plants expressing either Nkd1∆EAR or Nkd1SRDX were not different from plants expressing mCherry (Fig 6c). Taking all experiments together, the fact that Nkd1EARm and Nkd1ΔEAR complement the virulence defect of the ∆*nkd1* mutant but do not promote susceptibility of *A. thaliana* to Pst indicates that *U. maydis* uses other effectors to target TPL/TPRs in addition to Nkd1. Furthermore, the contribution of Nkd1 to the virulence of *U. maydis* likely depends on the interaction with targets other than TPL/TPRs. On the other hand, the fact that Nkd1SRDX is not able to complement the ∆*nkd1* mutant nor does it promote susceptibility of *A. thaliana* to Pst indicates that “mild” binding to TPL/TPRs (as seen for the WT Nkd1) promotes pathogen susceptibility whereas increased binding strength/specificity (as seen for the Nkd1SRDX, or by dramatically increasing nkd1 expression through the *cmu1* promoter) triggers resistance reactions independent of the suppression of PAMP-triggered ROS burst.

To assess the factors that could explain the lack of susceptibility to Pst in *A. thaliana* plants expressing Nkd1SRDX, we searched our RNAseq data for genes differentially expressed between these plants and those expressing the WT Nkd1. Clustering analysis indicated that plants expressing Ndk1SRDX form a unique group, different from Nkd1, 24 h after induction of transgene expression (Fig S7a). Thus, we focused on this time point. Amongst the most variable DEGs in this group, we found an enrichment for *PRs, R* genes, and genes involved in phytoalexin synthesis or metabolism (Fig 6d, Supplementary Table 2, Fig S7b, Supplementary Table 3), which are typical (SA) mediated responses (Robert-Seilaniantz et al., 2011). *PR* expression in particular, was between 1 and 3 orders of magnitude higher in plants expressing Nkd1SRDX compared to Nkd1 (WT) (Supplementary Table 2). In addition, we observed cell death symptoms in adult *A. thaliana* plants expressing Nkd1 and Nkd1SRDX that developed 3 days after induction of transgene expression, but the symptoms were stronger in the lines expressing Nkd1SRDX and these plants did not recover one week after the treatment. Plants expressing Nkd1∆EAR or mCherry did not show any cell death symptoms (Fig S8a). In *N. benthamiana*, cell death symptoms (observed 8 dpi) were not distinguishable between Nkd1 and Nkd1SRDX. Expression of Nkd1EARm lead to chlorosis mostly, and minimal cell death near the infiltration point. Expression of Nkd1∆EAR lead only to a very mild chlorosis and no cell death, nearly indistinguishable from the mCherry control. (Fig S8b).

To verify whether these responses were also observed macroscopically, in maize plants upon the expression of the different effector variants, we employed the pFoMV vectors and assessed their effect on *PR* and *NLR/HR* gene expression. In general, Nkd1 expressing plants showed increased expression of *PRs* compared with Nkd1∆EAR expressing plants. In the case of *ZmPR2*, only plants expressing Nkd1SRDX showed increased expression compared to GFP (control) or Nkd1∆EAR expressing plants. For *ZmPR1* and *ZmPR5*, Nkd1SRDX induced *PR* expression was not different from Nkd1 (WT) induced *PR* expression, even though in the former plants, viral spread and effector protein accumulation was much lower (Fig 6e, Fig S5a, S5b).

Maize plants expressing Nkd1 and Nkd1SRDX showed mild cell death symptoms (15 dpi) but these were not distinguishable from each other. Plants expressing either Nkd1EARm, Nkd1∆EAR or GFP did not show any cell death symptoms (Fig S8c). In addition, we did not observe upregulation of NLR gene expression in maize plants expressing any of the effector variants compared to the GFP control (Fig S7c).

Taken together our results indicate that increased binding of Nkd1 to TPL/TPRs leads to a pronounced upregulation of SA responsive genes, including *PRs*, whose expression pattern seems to be more conserved between monocot and dicot plants, and NLRs. Together, they increase resistance towards biotrophic pathogens despite the reduction in PAMP-triggered ROS burst.

## Discussion

Auxin has long been implicated in suppression of plant defenses since many pathogens produce this hormone and, in some cases, auxin biosynthetic genes are located in pathogenicity islands (Yamada, 1993). Besides the inhibition of SA mediated defenses, it is not clear how elevated auxin levels or signaling can promote disease susceptibility. Here we provide evidence that auxin signaling inhibits PAMP-triggered ROS burst, one of the earliest plant defense responses induced after microbial perception, and that this mechanism is exploited by the fungal effector Nkd1. We have shown that Nkd1 is secreted from *U. maydis* and translocated into the host nucleus where it targets TPL/TPRs. EAR motif-mediated binding of Nkd1 to TPL/TPRs prevents its recruitment to Aux/IAA transcriptional repressors, leading to elevated auxin signaling and subsequent inhibition of PTI. This inhibition is likely post-transcriptional, as we observed upregulation of many PTI-related genes in plants expressing Nkd1, amongst them *NADPH* oxidases. Nkd1 likely also promotes host susceptibility through the upregulation of jasmonic acid signaling, a well-documented phenomenon (Robert-Seilaniantz et al., 2011, Jiang et al., 2013). Therefore, our data suggests, that the main consequence of the upregulation of hormonal signaling induced by Nkd1 is defense inhibition rather than induction of gall development. Our results thus complement the previous observation that flg22 inhibits auxin signaling (Navarro et al., 2006, Navarro et al., 2004), they shed light on the mechanisms underlying the mutual antagonism between auxin signaling and PTI and why many pathogens (including non-gall inducing pathogens) manipulate host auxin signaling.

Previously, effector translocation into host cells was demonstrated using reporter strains or differential cultivar sets that rely on the hypersensitive response to indicate the presence of an effector in the plant cytosol (Mudgett et al., 2000, Wang et al., 2017). None of these tools are available in the *U. maydis*-maize pathosystem, so effector translocation has been demonstrated by technically demanding assays like immunolabeling followed by electron microscopy (Djamei et al., 2011, Redkar et al., 2015), functional read outs (Tanaka et al., 2014) or transgenic labeling approaches (Lo Presti et al., 2017). Here we have made use of the virulence defect of the *Δnkd1* mutant and the strong nuclear localization of Nkd1_(24-516)_ *in planta* to show that it is a translocated protein. Restricting Nkd1 nuclear entry (by fusion to the NES signal) prevents complementation of the *Δnkd1* virulence defect. This is consistent with the fact that Nkd1 expressed without signal peptide in maize cells is solely nuclear localized, that it interacts with TPL/TPRs (nuclear proteins) in the native system during infection, and with the fact that expression of WT Nkd1 in monocot and dicot plants leads to decreased PAMP-triggered ROS burst in an EAR motif-dependent manner, phenocopying the *tpl* knock out. Our results are thus in line with previous observations showing that knocking out *TPL/TPRs* in *A. thaliana* leads to increased susceptibility to Pst while overexpression of TPR1 leads to increased resistance to this pathogen (Zhu et al., 2010, Niu et al., 2019).

Even though WT Nkd1 inhibits PTI, this is unlikely to be the reason for the virulence defect observed in the *Δnkd1* mutant. PTI inhibiting effectors are highly redundant in *U. maydis*, as we have recently shown (Navarrete et al., 2019). We also identified another *U. maydis* effector, Jsi1, that targets TPL/TPRs. Jsi1 lacks any homology with Nkd1 but has a DLNxxP-type EAR motif that mediates its interaction with one of the C-terminal WD40 domains of TPL/TPRs. Deletion of *jsi1* does not lead to virulence defects and its expression *in planta* induces an overlapping but different set of hormone-responsive genes, predominantly from the jasmonate/ethylene branch (Darino et al., 2021). Therefore, the different domains of TPL/TPR proteins seem to control different hormone-responsive gene sets.

Despite the fact that Nkd1-mediated PTI inhibition depends on its interaction with TPL/TPRs via the EAR motif, complementation of the *Δnkd1* mutant with constructs encoding Nkd1 versions that do not bind to TPL/TPRs restored virulence to levels similar to WT. Since complementation of the *Δnkd1* mutant depends on the nuclear localization of the protein in the host cell, this indicates that the *Δnkd1* virulence defect is due to a second, yet unknown function of Nkd1 in the host cell nucleus. This scenario is feasible considering that Nkd1 is relatively large for an effector (65 kDa) but its ability to inhibit PTI depends on the very short C-terminal EAR motif, and that many effectors target several un-related host proteins (Xin et al., 2018, Park et al., 2012, Shi X. et al., 2018). Besides a second unidentified host target, another explanation for the virulence function of Nkd1 could be its interaction with one or more additional effectors which depend on Nkd1 to perform their function in maize.

Systematic Y2H screens between *U. maydis* effectors indicate a massive effector interactome (Alcantara et al., 2019) and effector-effector interactions have been shown to impact virulence in other pathosystems (Tzfira et al., 2004, Magori and Citovsky, 2011, Cain et al., 2008). Even though we could not detect an interaction between Nkd1 and Jsi1 (Alcantara et al., 2019), Nkd1 could potentially interact with other *U. maydis* effectors during maize infection. Therefore, analysis of Nkd1 interaction partners (plant or fungal) by IP-MS from infected maize tissue could help identify its second, non-redundant function.

Recent data suggest that TPL/TPRs may be involved in SA signaling as well. NPR3 and NPR4 are negative regulators of this defense pathway that repress SA responsive genes in the absence of a SA stimulus. Both, NPR3/4 have EAR motives and mutation of this motif abolishes the repressor activity of NPR4 (Ding et al., 2018). Further, NPR3 has recently been shown to interact with TPL (Altmann et al., 2020). Even more, another set of negative regulators of SA signaling, NIMINs, carry EAR motives and NIMIN1 has been shown to interact with TPL (Weigel et al., 2001, Consortium, 2011). However, the functional relevance of the interaction between these negative regulators of SA and TPL/TPRs has not been characterized so far. Thus, similar to what was observed here for the interaction between TPL/TPRs and AUX/IAAs, it is feasible that that Nkd1 and Nkd1SRDX could interfere with the interaction between TPL/TPRs and NIMINs or NPR3/4 but that the latter effector does it to a greater extent, for example by interfering with both, NIMINs and NPR3/4, whereas Nkd1 (WT) would interfere with a single of these systems. This is consistent with the fact that there is upregulation of SA-responsive genes upon expression of both effectors but Nkd1SRDX induces a far greater expression of SA-responsive genes like PRs and NLRs and that it does not induce pathogen susceptibility. What is more, Nkd1SRDX leads to a very marked reduction in the virulence of *U. maydis* and in the spread of FoMV in maize plants. This resistance response induced by Nkd1SRDX might involve the interaction between TPL/TPRs and R proteins. A recent report showed, that the interaction between AtTPR1 and the R protein SNC1 leads to increased expression of R genes and growth inhibition (Cai et al., 2019), similar to what we observed here in *A. thaliana*. Therefore, there seems to be an optimum strength for the EAR-mediated Nkd1 interaction with TPL/TPRs. In the WT Nkd1, its EAR motif mediates the interaction with TPL/TPRs to upregulate auxin signaling and inhibit PAMP triggered ROS burst. However, if the binding strength is increased and or the specificity for different TPL homologs is expanded (as in the case of Nkd1SRDX), or the effector expression is increased to non-physiological levels (*cmu1* promoter), additional defense responses are triggered that prevent pathogen susceptibility despite the fact that PAMP triggered ROS burst is inhibited.

In conclusion, we have identified a regulatory circuit in which a pathogen’s effector targets TPL/TPRs to upregulate auxin signaling and suppress PTI. Nonetheless, this immune suppression is not effective when other defense responses (mediated by PR and *R* genes) are triggered. This is in line with our experiments showing that NAA can inhibit PTI at a concentration optimum of 100 nM but not higher. It is also in line with previous observations (Navarro et al., 2006) where auxin-mediated promotion of Pst susceptibility was not effective when R gene-mediated responses were activated. Our finding that the EAR-motif is conserved among most Nkd1 orthologs, as well as of other TPL-binding effectors in *U. maydis* and other pathogens (Darino et al., 2019, Petre et al., 2015, Gawehns, 2014), suggests that fungi widely exploit the antagonism between growth and defense by targeting the central co-repressor TOPLESS. The discovery of the post-transcriptional mechanism through which TOPLESS-dependent elevated auxin signaling inhibits PTI awaits further research.

## Materials and methods

### Gene accession numbers

Umnkd1: UMAG_02299, Phnkd1: PHSY_006360, Srnkd1: sr13496, Sgnkd1: EX895_003371, Ssnkd1: SPSC_01711, Mpnkd1: BN887_03254. ZmTPL1: Zm00001d028481, ZmTPL2: Zm00001d040279, ZmTPL3: Zm00001d024523, ZmTPL4: Zm00001d047897, AtTPL: AT1G15750, AtTPR1: AT1G80490, AtTPR2: AT1G04130, AtTPR3: AT5G27030, AtTPR4: AT3G15880, NbTPR1: Niben101Scf19386g00002.1, NbTPR3: Niben101Scf00369g24005.1, MpTPR: Mapoly0051s0078.1, ZmIAA5: Zm00001d039624.

### Protein sequence analysis

For candidate effector genes, sequences were screened for the presence of a secretion signal with SignalP 5.0 (Almagro Armenteros et al., 2019). Protein alignments and (neighbor joining) phylogenetic trees were performed in CLC Main Workbench 7.7.2.

### Plasmids, cloning procedures, and generation of *U. maydis* strains

All plasmids were generated by standard molecular procedures (Sambrook et al., 2006) in *E. coli* Mach1 (Thermo Fisher Scientific, Waltham, MS, USA) was used for all DNA manipulations. All other plasmids were generated by the GreenGate system (Lampropoulos et al., 2013). The modules used were either amplified by PCR or obtained from the published system. Additionally, we generated two GreenGate destination vectors, pADGG and pBDGG, based on the pGAD and pGBKT7 backbone respectively, which were used for Y2H assays. Plasmids used for expression of VIGS assays in *N. benthamiana* were generated by Gateway Cloning (Katzen, 2007). For transient virus-mediated overexpression, PV101 (Bouton et al., 2018) was used to generate pFoMV:p19-P2A-mCherry-P2A-effector-myc and pFoMV:p19-P2A-mCherry-P2A-gfp-myc using NotI-XbaI cloning sites (p19: silencing suppressor, P2A: viral ribosome skipping motif). Unless otherwise stated, effector proteins were expressed without their secretion signals.

*U. maydis* knock out strains were generated by homologous recombination with PCR-derived constructs (Kämper, 2004). For complementation and protein expression, strains were generated by insertion of p123 derivatives into the *ip* locus (Loubradou, 2001). At least two independent strains were tested for each complementation construct. Transformants were verified by southern blot or PCR. All plasmids and strains used in this study can be found in the supplementary materials.

### Maize infection assays

Virulence assays and disease symptom scoring were performed as described by (Kämper et al., 2006). Briefly, *U. maydis* SG200 and its derivatives were cultured in liquid YepsLight (0.4% yeast extract, 0.4% peptone, and 2% sucrose) at 28°C to an OD_600nm_ of 0.6-0.8. Cells were pelleted by centrifugation at 2400 x g for 10 m and resuspended in H_2_O to an OD_600nm_ of 1. The suspensions were then syringe-inoculated into seven-day old B73 maize seedlings. Maize was grown in a temperature-controlled glasshouse (14 h light/10 h dark, 28°C/20°C) and disease symptoms were assessed 7 dpi. Filamentous growth of the *U. maydis* strains was tested by spotting the respective strains on CM agar (Brachmann et al., 2001) containing 1% activated charcoal. The experiments were repeated at least 3 times.

### Viral-mediated protein expression and biolistic bombardment in maize

Biolistic bombardment was performed according to (Djamei et al., 2011) with minor modifications. Briefly, 1.6 µm gold particles were coated with plasmid DNA encoding the indicated constructs under the control of the CaMV35S promoter. Bombardment was performed on seven-day old maize leaves cv B73 using a PDS-1000/HeTM instrument (BioRad) at 900 p.s.i. in a 27 Hg vacuum. Fluorescence was observed by confocal microscopy 18-24 h after bombardment. The experiments were repeated at least 3 times. Viral-mediated protein expression derived from the FoMV infectious clone was described earlier (Bouton et al., 2018). Plasmids were bombarded in whole-seedlings and plants were returned to the growth chamber.

### Analysis of gene expression in maize by qRT-PCR

Nine days post-bombardment, samples were taken and mCherry fluorescence was monitored as an indication of viral spread. Areas with positive mCherry fluorescence were used for RNA extraction. Total RNA was extracted with RNeasy Plant Mini Kit according to the manufacturer instructions (Qiagen, Cat. No. 74904). On-column DNase treatment was performed with RNase-Free DNase Set (Qiagen, Cat. No. 79254). 1 µg total RNA was used for cDNA synthesis with RevertAid™ Reverse Transcriptase (Thermo Fisher, Cat. No. EP0442). qPCRs were performed with GoTaq® qPCR mix (NEB, Cat. No. A6001) according to manufacturer’s instructions, using 0.5 µl of cDNA product. Relative amounts of amplicons were calculated according to the 2-∆∆Ct method (Livak and Schmittgen, 2001) with *CDK* (GRMZM2G149286/ Zm00001d010476) as reference gene (Lin et al., 2014).

### Cell death assessment in maize

Whole maize leaves were used to detect dead tissue with Trypan Blue staining (Fernanddez-Bautista et al., 2016). Leaves were completely immersed in a fresh Trypan Blue solution (containing: 85% lactic acid, 99% glycerol, phenol, ddH20 and Trypan Blue). After 30 minutes, stained leaves were immediately washed in 98% ethanol and left in it overnight. Ethanol solution was then replaced until the green tissue was cleared.

### Quantification of DR5 reporter activity in maize by epi-fluorescence microscopy

DR5:GFP was co-expressed with 35S:Nkd1-mCherry-3xHA or 35S:mCherry in maize leaves as described above and GFP fluorescence (Ex 480/40 nm Em 535/50 nm) was quantified from the double transformed cells using a widefield microscope equipped with transmitted and fluorescent light sources (Axio Imager.Z2). Briefly, the fluorescence intensity (535/50 nm) of each sample was calculated as the pixel intensity of the double transformed cells minus the pixel intensity of the un-transformed cells (background) using FIJI (Schindelin et al., 2012). At least 20 doubly transformed cells/treatment were analyzed per experiment. Experiments were repeated 3 times.

### Confocal microscopy

Confocal microscopy was performed with Zeiss LSM 700 or LSM 780 confocal microscope. GFP was excited at 488 nm using an argon laser. Fluorescence emission was collected between 500-540 nm. mCherry was exited at 561 nm and emission was collected between 578-648 nm. Images were processed with ZEN blue 2.3 lite. Fungal proliferation in infected tissue was assessed 5 dpi. Chitin was stained with wheatgerm agglutinin (WGA) coupled to AlexaFluor488 (Invitrogen). Plant cell walls and membrane compromised cells were stained with propidium iodide (Sigma‐Aldrich, St. Louis, MO, USA). Leaf samples were incubated with 10 µg/mL WGA‐AF488, 1 µg/mL propidium iodide and observed as described above. WGA‐ AF488 and propidium iodide excitation/detection condition were performed as for GFP and mCherry respectively.

### Protein secretion in *U. maydis*

For visualization of secreted proteins in infected tissues, maize plants were harvested 6-7 dpi and analyzed by confocal microscopy as described above. For plasmolysis experiments, infected maize tissue was infiltrated with 1 M mannitol and incubated at least 30 min prior to observation. Experiments were repeated at least 3 times.

Secreted proteins from fungal cultures were detected as previously (Djamei et al., 2011). *U. maydis* was grown in CM medium to an OD_600nm_ of 0.6-0.8, centrifuged, resuspended in AM medium and incubated for 5-6 h to induce filamentation. Cultures were centrifuged; supernatant proteins were precipitated with 10% trichloroacetic acid and 0.02% sodium deoxycholate and resuspended in 100 mM Tris pH 8. Proteins from the cell pellets were extracted by adding SDS loading buffer, a spatula tip of glass beads and vortexing for 10 min. All extracts were subjected to immunoblotting using α-HA (Sigma-Aldrich, St. Louis, MO, USA) for detection of effector proteins and α-Actin antibodies (Invitrogen, Waltham, MA, USA) for lysis control. Experiments were repeated 2 times.

### Yeast transformation and two-hybrid assays

All yeast protocols were performed according to the Yeast Protocols Handbook (Clontech, Mountainview, CA) with minor modifications. Strain AH109 was transformed with bait vectors (pGBKT7 and derivatives) and strain Y187 was transformed with prey vectors (pAD, pADGG and derivatives) by the LiAc/PEG method. All positive clones were verified for the presence of the corresponding plasmid by DNA extraction with 20 mM NaOH and PCR (Supplementary materials). For one to one matings, AH109 pBDGG-ZmTPL1, pBDGG-ZmTPL2, pBDGG-ZmTPL3, pBDGG-ZmTPL4, pBDGG-AtTPL, pBDGG-AtTPR1, AtTPR2, AtTPR4, or pBDGG-YFP were mated against Y187 pADGG-Nkd1, pADGG-Nkd1EARm, pADGG-Nkd1ΔEAR, pADGG-Nkd1SRDX, or pADGG-mCherry. Diploid cells were selected on SD –leu, -trp. For plate drop out assays, diploid cells were grown in liquid SD -leu, -trp o.n. Cells were pelleted by centrifugation at 500 x g for 3 min and resuspended in sterile H_2_O to an OD_600nm_ of 1. Serial dilutions were made in H_2_O and 5 µl of the suspensions were plated in SD -leu, -trp (growth control), SD -leu, -trp -his (intermediate stringency) and SD -leu, -trp, -his, -ade (high stringency). Growth in intermediate or high stringency media 4 days post inoculation indicated positive interactions. Experiments were repeated 3 times.

### Plant growth conditions and chemical treatments

*N. benthamiana* and *A. thaliana* plants were grown in controlled short-day conditions (8 h light/16 h dark, 21°C) in a soil:perlite mixture (4:1). Plants were watered by flooding for 15 min every two days. *A. thaliana* knock out and yucca OE lines were described previously (Supplementary materials). Estradiol inducible effector/mCherry lines were created by floral dipping using the following constructs: XVE:effector-3xMyc (Supplementary materials). All *A. thaliana* lines were stratified for 3 days at 4°C in the dark prior to sowing. Protein expression was induced by estradiol infiltration (20 µM, reporter and ROS burst assays) or spray (5µM, *Pseudomonas* infection assays). For auxin treatments, leaves of the appropriate age were infiltrated with 1-NAA (1-naphthaleneacetic acid, Duchefa) using a needleless syringe, plants were returned to the growth chamber and incubated for 6 h, and the leaf tissue was then processed as necessary.

### Protein production in *N. benthamiana* and Co-immunoprecipitation

For *in vivo* Co-IP assays, *A. tumefaciens* GV3101 (pSoup) carrying the expression constructs were grown o.n. in LB supplemented with the appropriate antibiotics at 28°C, centrifuged for 10 min at 1600 x g, resuspended in ARM buffer (Agrobacterium resuspension medium, 10 mM MES-NaOH pH 5.6, 10 mM MgCl_2_, 150 µM acetosyringone) to an OD_600nm_ of 0.2 and incubated for 3 h at RT. Cultures carrying the appropriate constructs were then mixed 1:1 and infiltrated in *N. benthamiana* with a needleless syringe. Plants were incubated for 48 h, frozen in liquid nitrogen, and total proteins were extracted from 300-450 mg of tissue in 2 ml IP buffer (HEPES 50 mM pH7.58, NaCl 50 mM, Glycerol 10%, EDTA 1 mM, Triton X-100 0.05%, PMSF 1 mM and 1 protease inhibitor tablet (Roche cOmplete EDTA free cat# 05056489001)/50 ml). Extracts were cleared by centrifugation 10 min at 20000 x g 3 times. Proteins were immunoprecipitated by adding 30 µl of α-c-Myc magnetic beads (µMACS Anti-c-myc Miltenyi Biotec, cat# 130-091-284) and incubated for 2 h at 4°C with rotation. Samples were washed 4 times with 300 µl IP buffer and proteins were eluted by adding 50 µl of 2x SDS loading buffer at 95°C. 10-15 μl of the extracts were analyzed by SDS-PAGE followed by western blot with α-c-Myc (Sigma-Aldrich, St. Louis, MO, USA) or α-HA (Sigma-Aldrich, St. Louis, MO, USA) antibodies. Experiments were repeated at least 3 times.

For Co-IP assays from maize tissues, one-week old plants were infected with *U. maydis* carrying the appropriate constructs (Supplementary materials). Plants were incubated for 7 days, frozen in liquid nitrogen and total proteins were extracted from 2-3 g of tissue in 25 ml of IP buffer. Proteins were immunoprecipitated by adding 150 ul of α-c-HA magnetic beads (µMACS Anti-c-myc Miltenyi Biotec, cat# 130-094-255) as described above. 10-25 μl of the extracts were analyzed by SDS-PAGE followed by western blot with α-HA or α-TPL (Eurogentec, Belgium). The latter (polyclonal) antibodies were developed in rabbit against the peptide CNEQLSKYGDTKSAR, that is conserved across TPL/TPR proteins (Darino et al., 2019). Experiments were repeated 2 times.

### VIGS in *N. benthamiana*

Three-week old *N. benthamiana* plants were infiltrated with a mixture of *A. tumefaciens* GV3101 (pSoup) strains carrying pTRV1 (OD_600nm_ 0.2), pTRV2-tpr1 (OD_600nm_ 0.1), and pTRV2-tpr3 (OD_600nm_ 0.1) in ARM. Control plants were infiltrated with a mixture of strains carrying pTRV1 (OD_600nm_ 0.2) and pTRV2-gfp (OD_600nm_ 0.2) (Ratcliff et al., 2001). Plants were then grown for 2-3 weeks before being analyzed. Experiments were repeated 3 times.

### DR5 reporter induction in *N. benthamiana* and *A. thaliana*

5-6-week-old *A. thaliana* or 4-5-week old *N. benthamiana* plants were used for the assay. *N. benthamiana* was infiltrated with *A. tumefaciens* carrying the appropriate constructs resuspended in ARM buffer to a final OD_600nm_ of 0.1 and incubated for 72 h. Leaf disks (4 mm) were cut, floated on water, and GFP fluorescence (485/528 nm) was assessed in a microplate reader (Synergy H1, BioTek). *A. thaliana* leaf disks were incubated o.n. prior to fluorescence measurement. At least two *N. benthamiana* or five *A. thaliana* plants per construct/genotype were used in each experiment. Experiments were repeated at least 3 times.

### ROS burst assays in *N. benthamiana, A. thaliana*, and maize

*A. thaliana* and *N. benthamiana* plants were grown as described for the reporter induction assays. Maize plants were 7 days old and only the first true leaf was used for the assay. Leaf disks (4 mm) were cut and floated on water o.n. For plants expressing pFoMV vectors, 8-9 days after bombardment areas with positive mCherry fluorescence (indicating viral spread) were used to cut leaf disks which were then floated on water o.n. Water was then removed, and elicitors were added. flg22 elicitation solution consisted of Horseradish peroxidase (HRP 10 µg/ml, Sigma-Aldrich cat# P6782), L-012 (34 µg/ml Fujifilm WAKO cat# 120-04891) and flg22 (100 nM) in H_2_O. Chitin elicitation solution was prepared as follows: 50 mg of chitin (Sigma-Aldrich cat# C9752) were ground with mortar and pestle in 5 ml of H_2_O for 5 min, transferred to a falcon tube, microwaved for 40 sec, sonicated for 5 min, centrifuged at 1800 x g for 5 min, the supernatant was transferred to a new tube, vortexed for 15 min, and stored at 4**°**C. Before use, the suspension was diluted 1:1 in H_2_O and supplemented with HRP (as described above) and luminol (34 µg/ml Sigma-Aldrich cat# 123072). Reactive oxygen species production was monitored by luminescence over 30-40 min in a microplate reader (Synergy H1, BioTek). At least 3 *N. benthamiana* or *A. thaliana* plants per construct/genotype were used in each experiment. Experiments were performed at least 3 times.

### Pseudomonas syringae infections in A. thaliana

Plants were grown for one month under 12 h light/12 h dark at 21°C, sprayed with 5 µM estradiol in 0.01% Silwet, and incubated o.n. *P. syringae* pv. tomato DC3000 was infiltrated in the leaf tissue with a needleless syringe (OD_600nm_ = 0.0002). Plants were covered with a plastic dome and returned to the growth chamber for 2 days. Leaf disks were cut, homogenized in 2 ml tubes containing MgCl_2_ and metal beads, and bacterial load was assessed by plate counts in King’s B media supplemented with Rifampicin (50 µg/ml). Plates were incubated at R.T. until colonies were visible. The experiment was repeated 3 times.

### RNAseq experiments

*A. thaliana* seedlings were grown vertically for 7 days in ½ MS, 2% sucrose plates containing a nylon mesh (100 µm pore size, SEFAR®). Light conditions were 16 hr/8 hr light/dark cycle at 21°C. Protein expression was induced by transferring the nylon mesh with the seedlings onto a new ½ MS plate supplemented with β-estradiol (5 µM). Seedlings were then returned to the growth chamber and incubated for a further 5 h or 24 h and frozen in liquid nitrogen. The experiment was repeated 3 times. 30 mg of tissue were used for RNA or protein extraction. Expression of β-estradiol inducible constructs was assessed by western blot with α-c-Myc antibodies as described earlier. RNA extractions were performed with RNeasy Plant kit (QIAGEN) and mRNA was isolated by Poly(A) RNA Selection kit (Lexogen) from 1 µg of total RNA. Library preparation was done using Lexogen SENSE kit. Libraries, single end 75, were sequenced in an Illumina NextSeq550 sequencer. Library preparation and sequencing was performed by the VBCF NGS Unit (www.viennabiocenter.org/facilities).

### RNAseq and differential gene expression analysis

Raw RNA sequencing data was aligned using STAR version 2.5.1 (Dobin et al., 2013). The average sequencing depth per sample was 33.5 million reads (st.dev. 3.7 mln reads).

Gene expression for all TAIR10 protein-coding genes was calculated using featureCounts tool from Subread (version 1.4.6) package (Liao et al., 2014). Expression heatmaps were created from normalized expression data (TPMs) averaged among 3 biological replicates using pheatmap R package (with scaling for each gene (scale=“row”).

Differential gene expression was accessed with DESeq2 (Love et al., 2014) using raw count tables generated by featureCounts. Each sample had 3 replicates.

### Statistical analyses

Maize infection assays were analyzed by the Fisher exact test in R, as described by (Stirnberg and Djamei, 2016). All other statistical analyses were performed with GraphPad Prism 8.0. ROS burst data and fluorescence measurements from DR5 reporter assays were analyzed by the Student’s t-test, Tukey’s ANOVA or Bonferoni’s two-way ANOVA. *P. syringae* infections (log CFU/cm^2^) was analyzed by Tukey’s ANOVA. Statistical significance was evaluated at the level of p<0.05. For differential gene expression from RNAseq data, threshold for calling a gene differentially expressed between two conditions or samples was taken to be: log2 fold change > 0.6 and adjusted p value <0.01.

## Data availability

The authors declare that all the data supporting the current findings are available within the paper, supplementary information file, source data file and publically available data bases (GEO Id for RNAseq data: GSE141597).

## Acknowledgements

The research leading to these results has received funding from the European Research Council under the European Union’s Seventh Framework Programme ERC-2013-STG, Grant Agreement: 335691, the Austrian Science Fund (FWF): [P27818-B22, I 3033-B22], the Austrian Academy of Sciences (OEAW) and the Deutsche Forschungsgemeinschaft (DFG, German Research Foundation) under Germany’s Excellence Strategy -EXC 2070-390732324.

We would like to thank the GMI/IMBA/IMP core facilities for excellent technical support, especially the BioOptics and Molecular Biology Services. We thank the Plant Sciences and Next Generation Sequencing Facilities at the Vienna BioCenter Core Facilities GmbH (VBCF). We are grateful to Jirí Friml and Jürgen Kleine-Vehn laboratories for providing useful *A. thaliana* lines. We thank Mathias Madalinski for peptide synthesis and to Dr. J. Matthew Watson for proofreading and valuable feedback on the manuscript.

## Author Contribution

Conceptualization: Fernando Navarrete, Armin Djamei.

Funding acquisition: Armin Djamei.

Investigation: Fernando Navarrete, Michelle Gallei, Martin A. Darino, Aleksandra Kornienko, Indira Saado, Khong-Sam Chia, Mamoona Khan.

Methodology: Fernando Navarrete, Armin Djamei.

Project administration: Fernando Navarrete.

Resources: Fernando Navarrete, Michelle Gallei, Aleksandra E. Kornienko, Martin A. Darino, Indira Saado, Janos Bindics, Armin Djamei.

Supervision: Armin Djamei.

Writing: Fernando Navarrete, Armin Djamei.

## Supplementary information

Supplementary Figure S1

Supplementary Figure S2

Supplementary Figure S3

Supplementary Figure S4

Supplementary Figure S5

Supplementary Figure S6

Supplementary Figure S7

Supplementary Figure S8

Supplementary Table 1. Genes differentially expressed in *A. thaliana* seedling expressing different effector constructs 5h post estradiol treatment. Linked to Fig 4d.

Supplementary Table 2. Genes differentially expressed in *A. thaliana* seedling expressing different effector constructs 24h post estradiol treatment. Linked to Fig 6d.

Supplementary Table 3. Genes differentially expressed in *A. thaliana* seedlings expressing different effector constructs 24h post estradiol treatment. Linked to Fig S7b.

Supplementary Materials. List of primers, plasmids, bacterial strains, *U. maydis* strains, plant lines.

