## Supplementary information for "TOPLESS promotes plant immunity by repressing auxin signaling and is targeted by the fungal effector Naked1"

Supplementary Figure S1……………………………………………………………………………..2

Supplementary Figure S2…………………………………………………………………..……3

Supplementary Figure S3……………………………………………………………………………..4

Supplementary Figure S4……………………………………………………………………………..5

Supplementary Figure S5……………………………………………………………………………..7

Supplementary Figure S6……………………………………………………………………………..9

Supplementary Figure S7……………………………………………………………………………10

Supplementary Figure S8…………………………………………………………………..…12

Supplementary Table 1. Genes differentially expressed in *A. thaliana* seedling expressing different effector constructs 5h post estradiol treatment. Linked to Fig 4d…………………….…14

Supplementary Table 2. Genes differentially expressed in *A. thaliana* seedling expressing different effector constructs 24h post estradiol treatment. Linked to Fig 6d…………………..…19

Supplementary Table 3. Genes differentially expressed in *A. thaliana* seedlings expressing different effector constructs 24h post estradiol treatment. Linked to Fig S7b……………………21

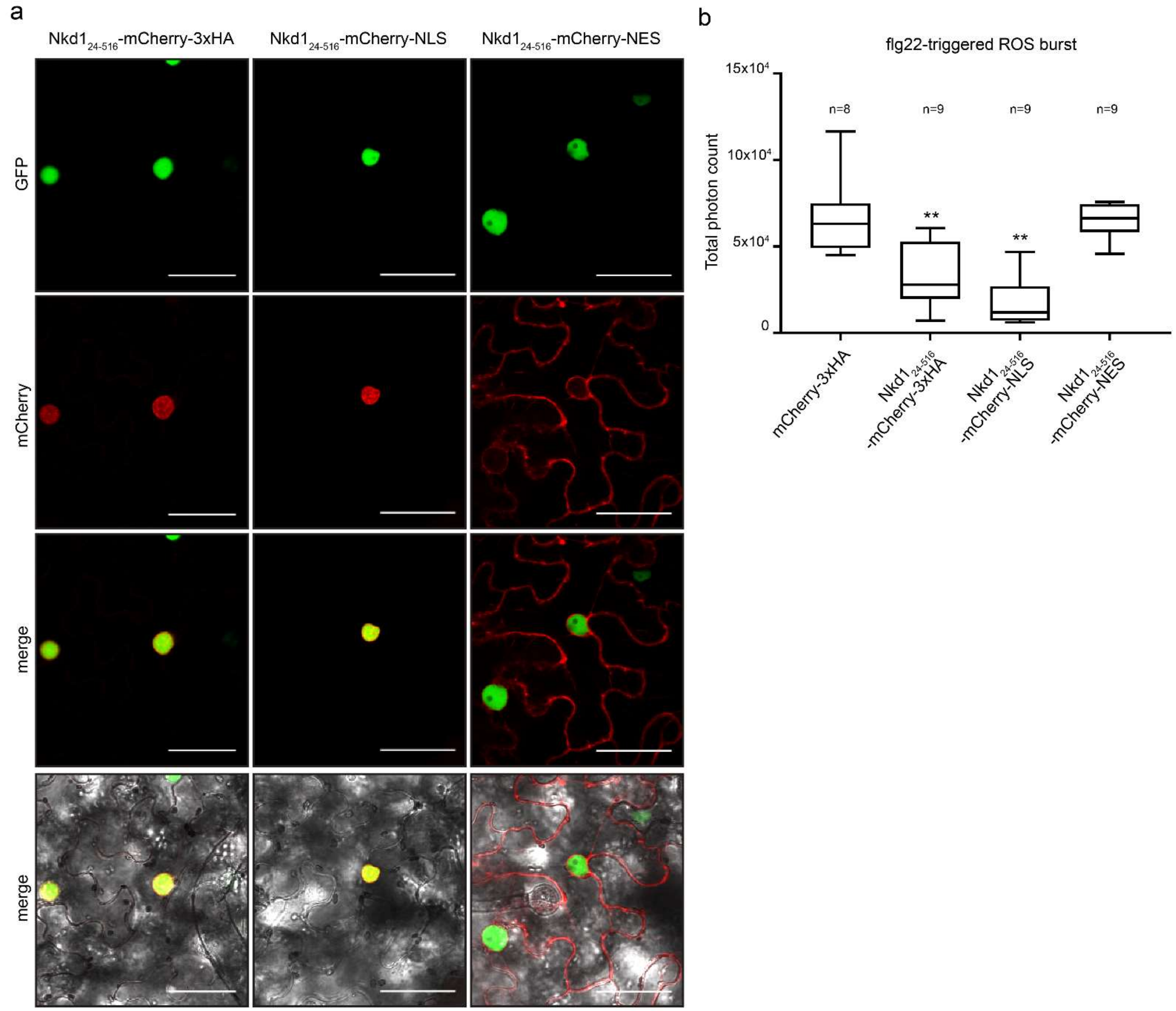

**Supplementary Figure S1.**

**a.** Localization of different Nkd1-mCherry fusions in the epidermis of *N. benthamiana*. Panels from top: GFP, mCherry, GFP-mCherry merge, bright field-GFP-mCherry merge. Scale bar = 50 µm. **b.** flg22-triggered ROS burst of plants expressing the constructs shown in a. Plants expressing mCherry were used as the positive control. Total photon counts over 40 minutes are shown as box plots. Data is a pool of three independent experiments, n= number of plants used for each group (* $p<0.05$, ** $p<0.01$, ANOVA, Tukeys).

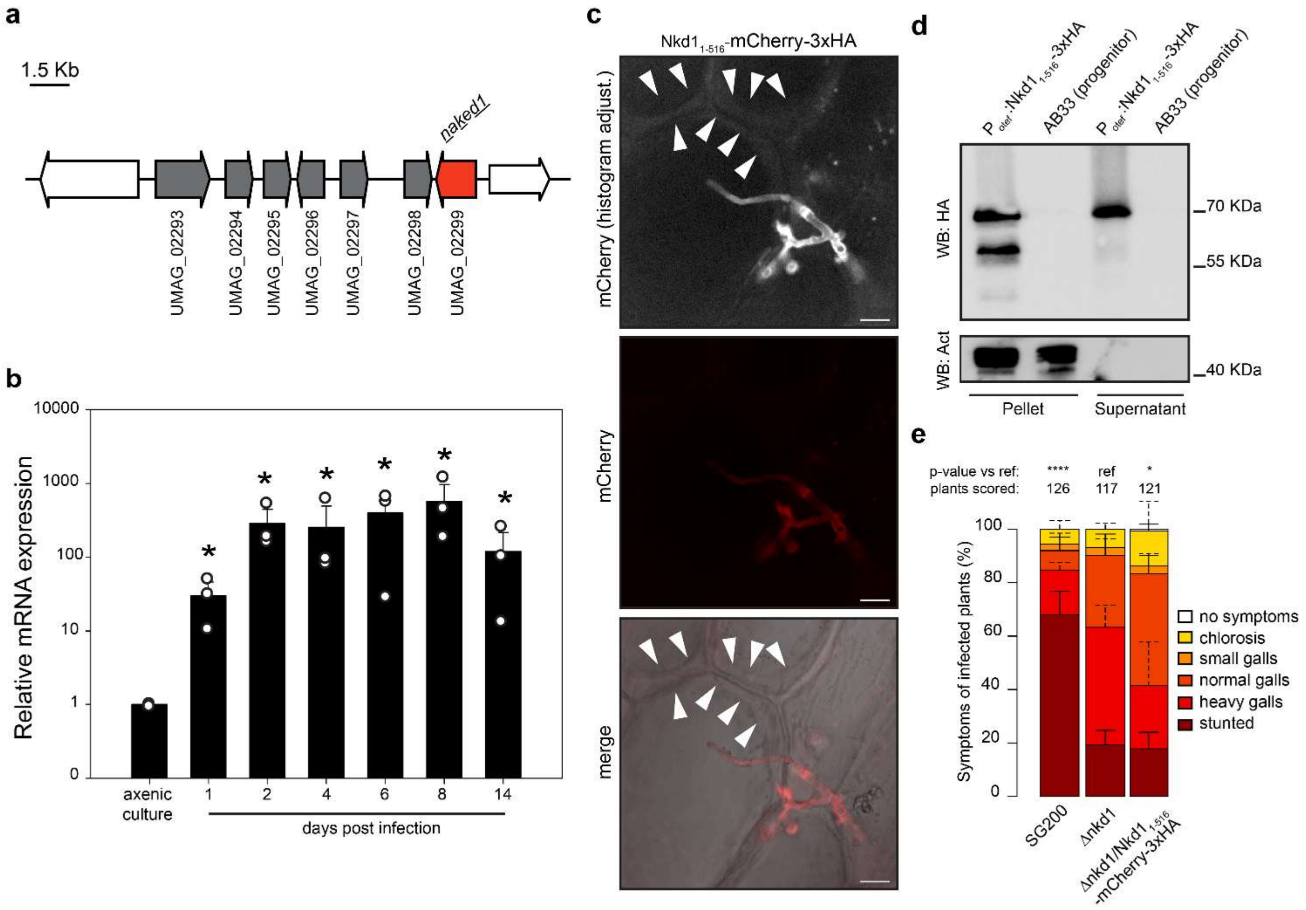

**Supplementary Figure S2**

**a.** Schematic representation of the *nkd1* in *U. maydis* effector cluster 5-21 on chromosome 5. *nkd1* is depicted in red. Genes in gray are part of the effector gene cluster 5-21 and contain a predicted secretion signal. Genes in white are the neighboring genes of the gene cluster and do not contain secretion signals. **b.** Transcriptional regulation of *nkd1* during *U. maydis* infection of maize. Expression was monitored by qPCR and normalized to the constitutively expressed gene *ppi*. Data is mean ± SD, n=3 (*p<0.05, ANOVA with Benjamini-Hochberg correction for multiple comparisons). **c.** Localization of Nkd1 during host colonization. Plants infected with a strain expressing Nkd1$_{1-516}$-mCherry-3xHA were plasmolyzed with 1 M mannitol and visualized by confocal microscopy. Upper panel: mCherry fluorescence (depicted in white) with histogram adjustment to highlight lower intensity pixels, middle panel: mCherry fluorescence (original), bottom panel: bright field-mCherry merge. Arrowheads indicate the plant plasma membrane and the apoplastic space in-between them. mCherry signal in the apoplastic space indicates secretion of soluble protein. Scale bar = 10 µm. **d.** Secretion of Nkd1 in axenic culture. Nkd1$_{1-516}$-3xHA was expressed in the strain AB33 driven by the strong, constitutive *otef* promoter. Total proteins were extracted from the pellet and secreted proteins were precipitated from the culture supernatant. The extracts were subjected to western blot with α-HA or α-Actin antibodies. Nkd1-3xHA could be detected in both, pellet and supernatant whereas the non-secreted control, actin, was only detected in the pellet fraction. **e.** Disease symptom scoring of maize seedlings infected with *U. maydis* (7 dpi). SG200 (progenitor strain), Δ*nkd1* and its complementation with Nkd1$_{1-516}$-mCherry-3xHA. Data represent mean ± SD from three independent experiments, n= total number of scored plants. Significant differences between strains were analyzed by Fisher's exact test with Benjamini-Hochberg correction for multiple comparisons (*p<0.05, ** p<0.01, *** p<0.001, **** p<0.0001).

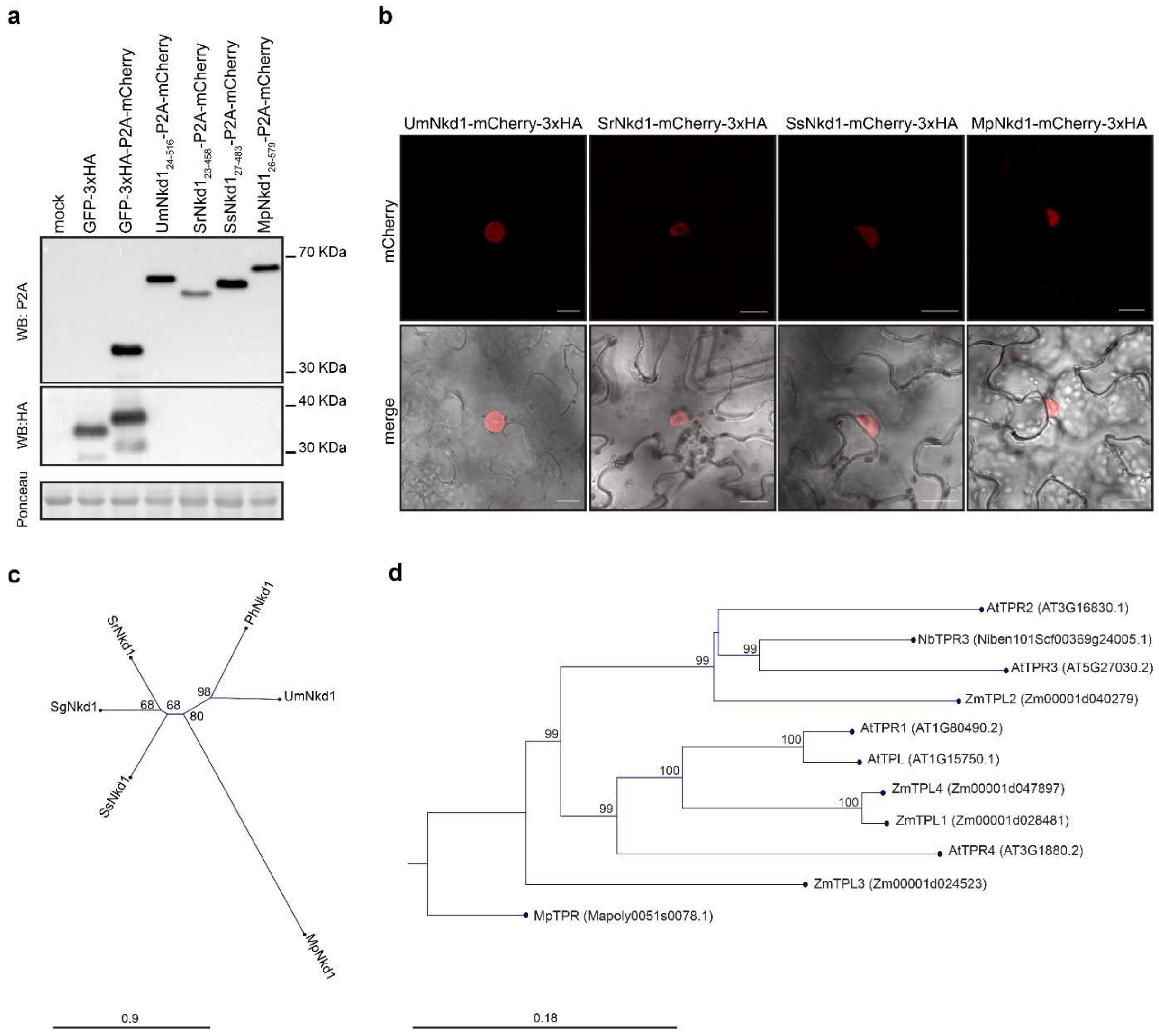

**Supplementary Figure S3.**

**a.** Expression of Nkd1 orthologs in *N. benthamiana* was assessed by western blot with α-P2A antibodies. Western blot with α-HA antibodies shows the specificity of the α-P2A antibodies. **b.** Localization of different Nkd1 orthologs in *N. benthamiana*. Orthologs were fused to mCherry-3xHA and localization was assessed by confocal microscopy. All orthologs localize to the plant nucleus. Scale bar = 20 µm. **c.** Maximum likelihood, un-rooted phylogenetic tree of Nkd1 orthologs. **d.** Maximum likelihood phylogenetic tree of TPL/TPR proteins shown in Fig 3. Mp*TPR* (*Marchantia polymorpha*) was used as the outgroup. For c and d, branch length represents genetic distance according to the Kimura 2-parameter.

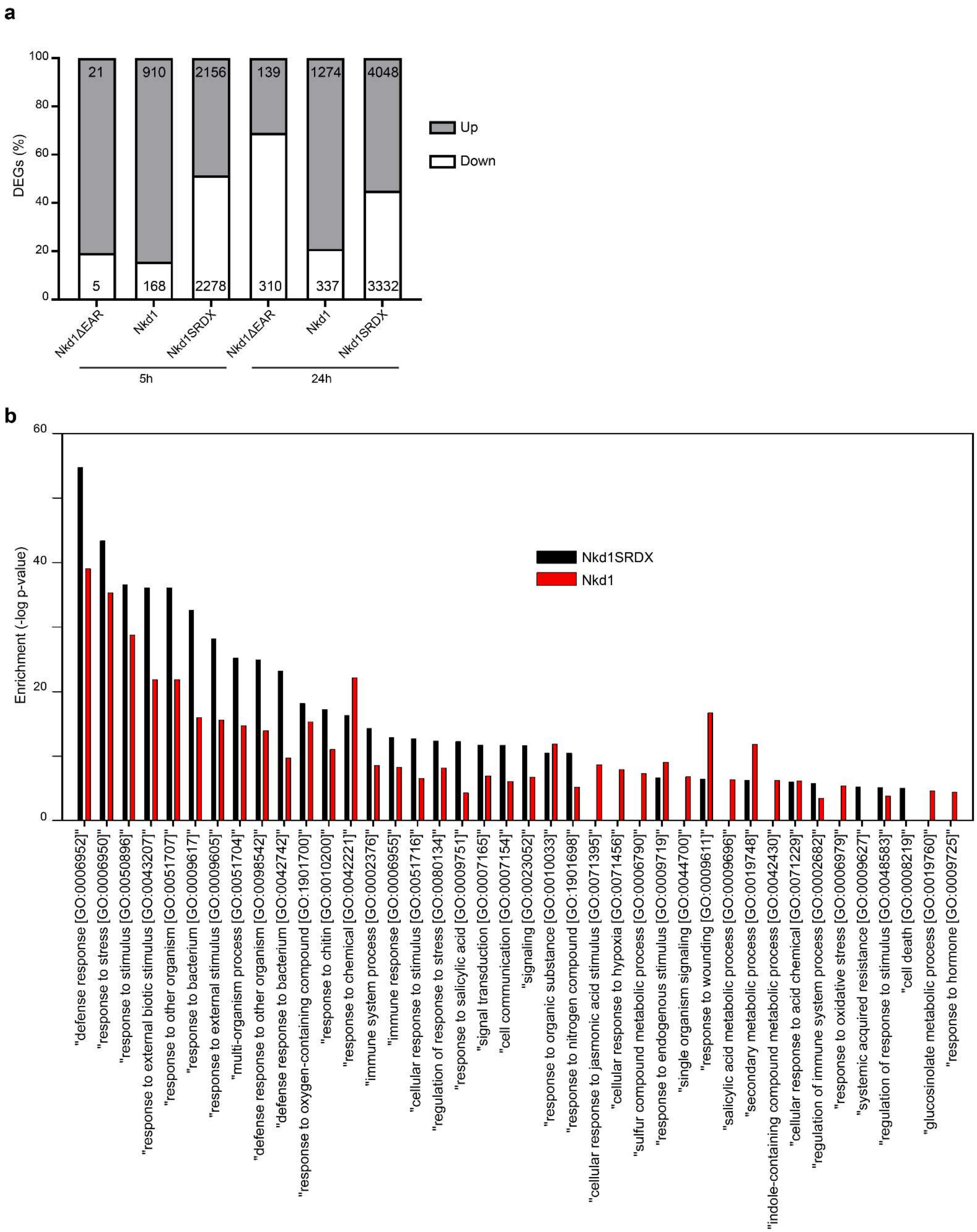
a
DEGs (%)
100
80
60
40
20
0
21
910
2156
139
1274
4048
5
168
2278
310
337
3332
Up
Down
Nkd1ΔEAR
Nkd1
Nkd1SRDX
Nkd1ΔEAR
Nkd1
Nkd1SRDX
5h
24h
b
Enrichment (-log p-value)
60
40
20
0
Nkd1SRDX
Nkd1
"defense response [GO:0006952]"
"response to stress [GO:0006950]"
"response to stimulus [GO:0050896]"
"response to external biotic stimulus [GO:0043207]"
"response to other organism [GO:0051707]"
"response to bacterium [GO:0009617]"
"response to external stimulus [GO:0009605]"
"multi-organism process [GO:0051704]"
"defense response to other organism [GO:0098542]"
"defense response to bacterium [GO:0042742]"
"response to oxygen-containing compound [GO:1901700]"
"response to chitin [GO:0010200]"
"response to chemical [GO:0042221]"
"immune system process [GO:0002376]"
"immune response [GO:0006955]"
"cellular response to stimulus [GO:0051716]"
"regulation of response to stress [GO:0080134]"
"response to salicylic acid [GO:0009751]"
"signal transduction [GO:0007165]"
"cell communication [GO:0007154]"
"signaling [GO:0023052]"
"response to organic substance [GO:0010033]"
"response to nitrogen compound [GO:1901698]"
"cellular response to jasmonic acid stimulus [GO:0071395]"
"cellular response to hypoxia [GO:0071456]"
"sulfur compound metabolic process [GO:0006790]"
"response to endogenous stimulus [GO:0009719]"
"single organism signaling [GO:0044700]"
"response to wounding [GO:0009611]"
"salicylic acid metabolic process [GO:0009696]"
"secondary metabolic process [GO:0019748]"
"indole-containing compound metabolic process [GO:0042430]"
"cellular response to acid chemical [GO:0071229]"
"regulation of immune system process [GO:0002682]"
"response to oxidative stress [GO:0006979]"
"systemic acquired resistance [GO:0009627]"
"regulation of response to stimulus [GO:0048583]"
"cell death [GO:0008219]"
"glucosinolate metabolic process [GO:0019760]"
"response to hormone [GO:0009725]"

**Supplementary Figure S4.**
**a.** Differentially expressed genes (DEGs, fold change > 1.5, p-value <0.01) in *A. thaliana* lines expressing Nkd1 and its EAR mutants. Plants expressing mCherry were used as the reference control. **b.** GO enrichment for biological function of genes upregulated in Nkd1 and Nkd1SRDX 5 h post induction of transgene expression.

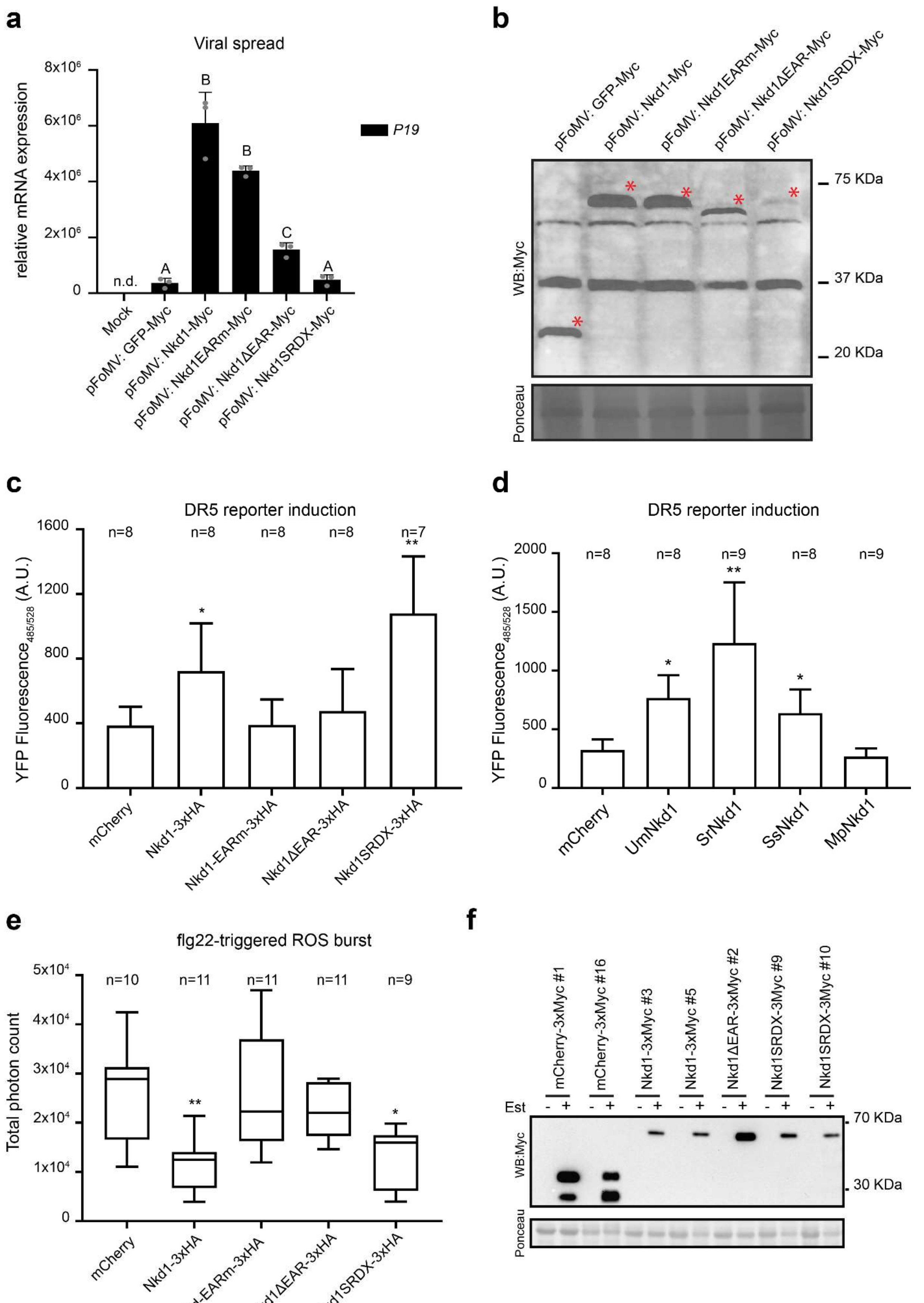

a
Viral spread
relative mRNA expression
P19
n.d.
Mock
pFoMV: GFP-Myc
pFoMV: Nkd1-Myc
pFoMV: Nkd1EARm-Myc
pFoMV: Nkd1ΔEAR-Myc
pFoMV: Nkd1SRDX-Myc
b
WB:Myc
75 KDa
37 KDa
20 KDa
Ponceau
c
DR5 reporter induction
YFP Fluorescence485/528 (A.U.)
mCherry
Nkd1-3xHA
Nkd1-EARm-3xHA
Nkd1ΔEAR-3xHA
Nkd1SRDX-3xHA
d
DR5 reporter induction
mCherry
UmNkd1
SrNkd1
SsNkd1
MpNkd1
e
flg22-triggered ROS burst
Total photon count
mCherry
Nkd1-3xHA
Nkd-EARm-3xHA
Nkd1ΔEAR-3xHA
Nkd1SRDX-3xHA
f
mCherry-3xMyc #1
mCherry-3xMyc #16
Nkd1-3xMyc #3
Nkd1-3xMyc #5
Nkd1ΔEAR-3xMyc #2
Nkd1SRDX-3Myc #9
Nkd1SRDX-3Myc #10
Est
WB:Myc
70 KDa
30 KDa
Ponceau

**Supplementary Figure S5.**
**a.** Spread of viral constructs expressing the different Nkd1 variants shown in Fig 5c. Expression of *P19* (encoded in the viral constructs) was monitored by qRT-PCR and normalized to Zm*CDK* 9dpi. Data is mean ± SD, n=3 (different letters indicate statistical difference, $p<0.05$, by Tukey's ANOVA, n.d.: not detected). **b.** expression of viral-encoded Nkd1 variants was assessed by western blot with α-Myc antibodies. (* full length protein). **c.** DR5:YFP reporter induction in *N. benthamiana* plants expressing Nkd1 or its EAR mutants. Data (mean ± SD) is is a pool of 3 independent experiments, n= number of plants used for each group. Constructs shown here are the same ones used in Figure 4c. **d.** DR5:YFP reporter induction in *N. benthamiana* plants expressing different Nkd1 orthologs. Constructs shown here are the same ones used in Figure 3b. For c and d, data (mean ± SD) is a pool of 4 independent experiments n= number of plants used for each group, (* $p<0.05$, ** $p<0.01$, Tukey's ANOVA). **e.** flg22 triggered ROS burst in *N. benthamiana* expressing Nkd1 or its EAR mutants. Plants expressing mCherry were used as the positive control. Total photon counts over 40 minutes are shown as box plots. Data is a pool of 3 independent experiments, n= number of plants used for each group (* $p<0.01$, ** $p<0.01$, Tukey's ANOVA). **f.** Expression of Nkd1 and its EAR mutants in *A. thaliana* 24h hours after β-estradiol treatment. Lines shown here are the same ones shown in Fig 5d and 5e.

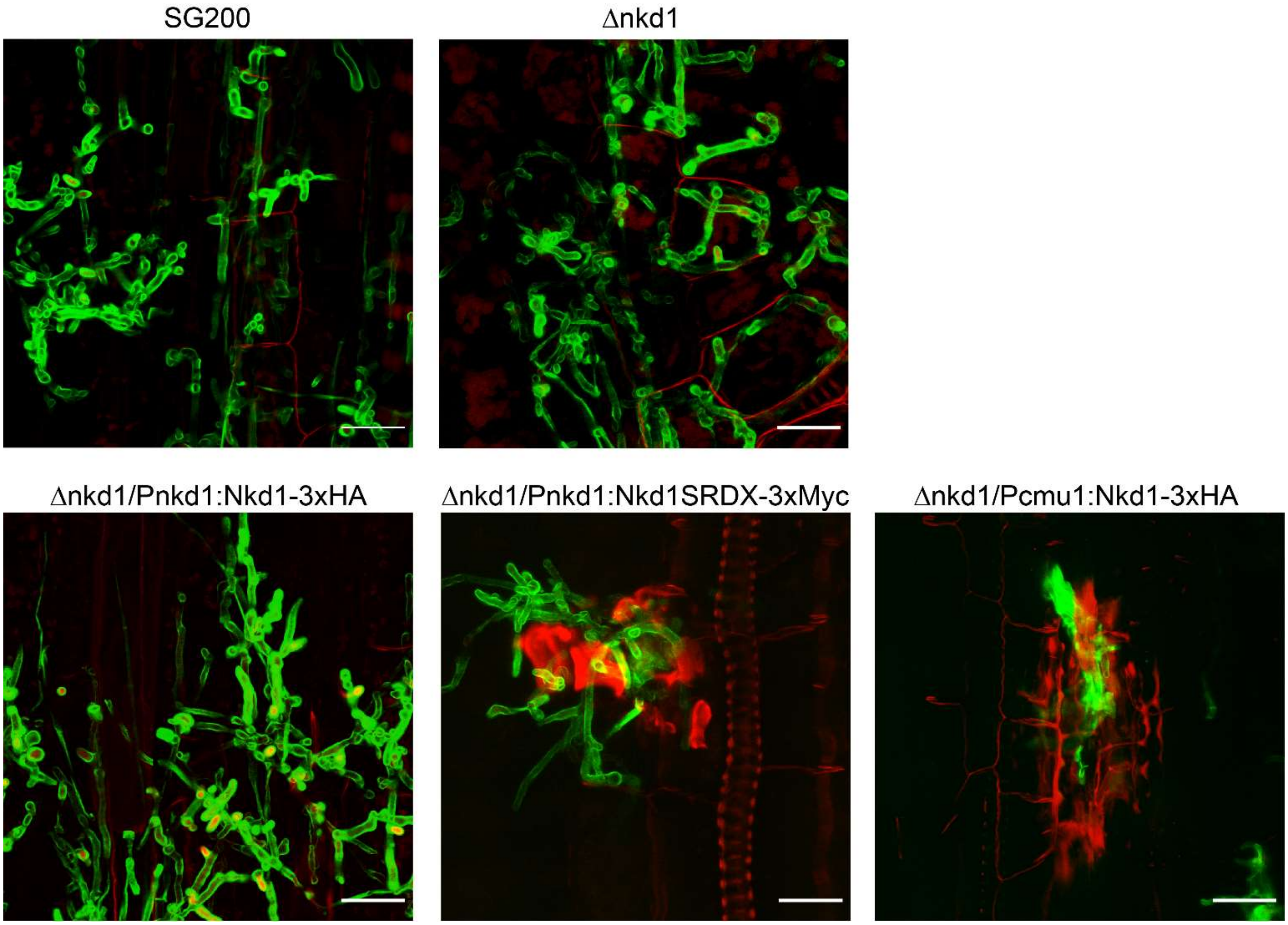

**Figure S6.**
Microscopic analysis of different *U. maydis* strains infecting maize. **a.** Plants infected with different *U. maydis* strains (5dpi) were stained with WGA-AlexaFluor488 (green) to visualize fungal chitin and propidium iodide (red) to observe dead cells and plant cell walls. Scale bars = 20 μm.

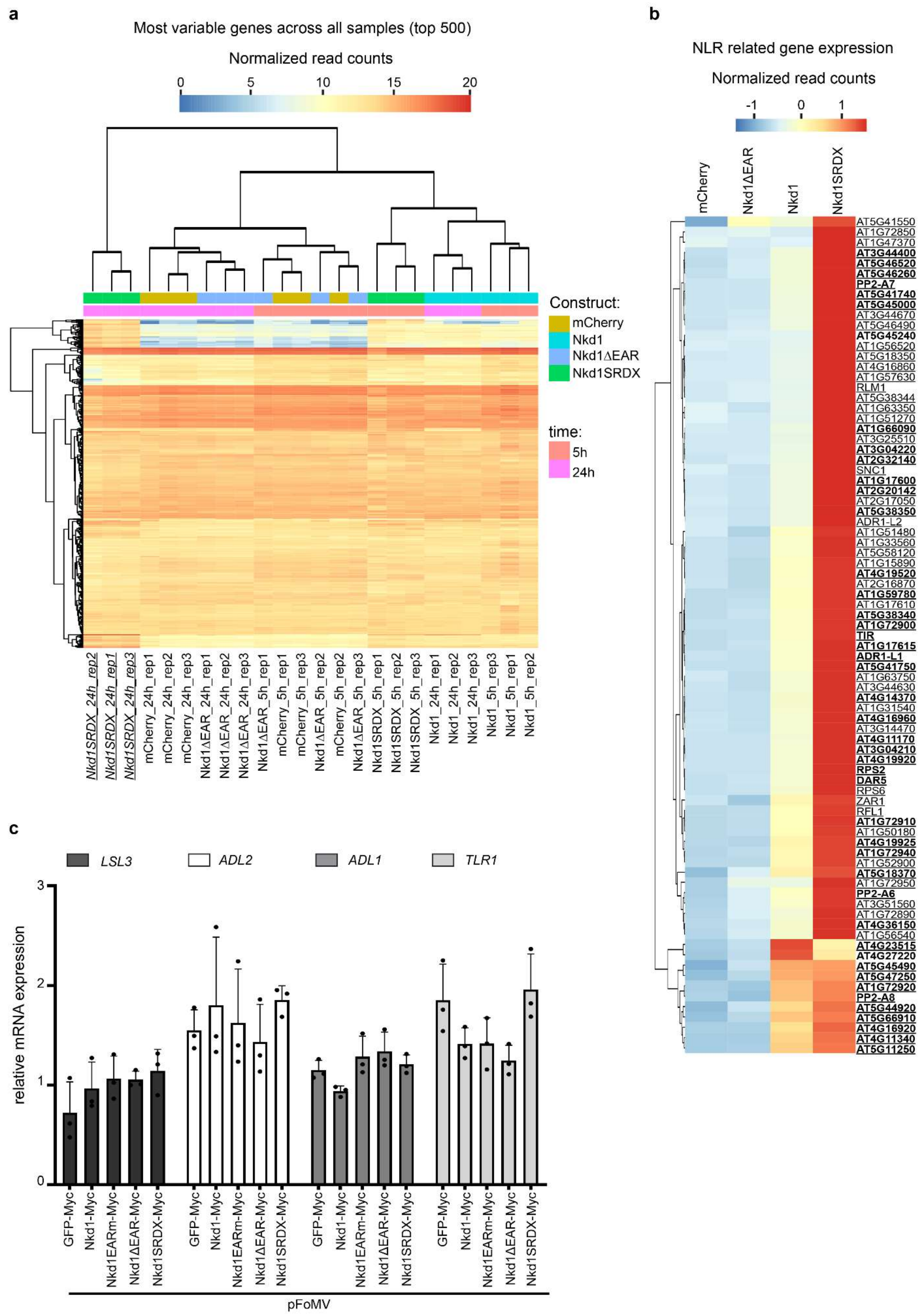
a
Most variable genes across all samples (top 500)
Normalized read counts
0
5
10
15
20
Construct:
mCherry
Nkd1
Nkd1ΔEAR
Nkd1SRDX
time:
5h
24h
Nkd1SRDX_24h_rep2
Nkd1SRDX_24h_rep1
Nkd1SRDX_24h_rep3
mCherry_24h_rep1
mCherry_24h_rep2
mCherry_24h_rep3
Nkd1ΔEAR_24h_rep1
Nkd1ΔEAR_24h_rep2
Nkd1ΔEAR_24h_rep3
Nkd1ΔEAR_5h_rep1
mCherry_5h_rep1
mCherry_5h_rep3
Nkd1ΔEAR_5h_rep2
mCherry_5h_rep2
Nkd1ΔEAR_5h_rep3
Nkd1SRDX_5h_rep1
Nkd1SRDX_5h_rep2
Nkd1SRDX_5h_rep3
Nkd1_24h_rep1
Nkd1_24h_rep2
Nkd1_24h_rep3
Nkd1_5h_rep3
Nkd1_5h_rep1
Nkd1_5h_rep2
b
NLR related gene expression
Normalized read counts
-1
0
1
mCherry
Nkd1ΔEAR
Nkd1
Nkd1SRDX
AT5G41550
AT1G72850
AT1G47370
AT3G44400
AT5G46520
AT5G46260
PP2-A7
AT5G41740
AT5G45000
AT3G44670
AT5G46490
AT5G45240
AT1G56520
AT5G18350
AT4G16860
AT1G57630
RLM1
AT5G38344
AT1G63350
AT1G51270
AT1G66090
AT3G25510
AT3G04220
AT2G32140
SNC1
AT1G17600
AT2G20142
AT2G17050
AT5G38350
ADR1-L2
AT1G51480
AT1G33560
AT5G58120
AT1G15890
AT4G19520
AT2G16870
AT1G59780
AT1G17610
AT5G38340
AT1G72900
TIR
AT1G17615
ADR1-L1
AT5G41750
AT1G63750
AT3G44630
AT4G14370
AT1G31540
AT4G16960
AT3G14470
AT4G11170
AT3G04210
AT4G19920
RPS2
DAR5
RPS6
ZAR1
RFL1
AT1G72910
AT1G50180
AT4G19925
AT1G72940
AT1G52900
AT5G18370
AT1G72950
PP2-A6
AT3G51560
AT1G72890
AT4G36150
AT1G56540
AT4G23515
AT4G27220
AT5G45490
AT5G47250
AT1G72920
PP2-A8
AT5G44920
AT5G66910
AT4G16920
AT4G11340
AT5G11250
c
LSL3
ADL2
ADL1
TLR1
relative mRNA expression
0
1
2
3
GFP-Myc
Nkd1-Myc
Nkd1EARm-Myc
Nkd1ΔEAR-Myc
Nkd1SRDX-Myc
pFoMV

**Supplementary Figure S7.**

**a.** Expression pattern (Euclidian clustering) of the top 500 most variably expressed genes across all *A. thaliana* seedlings. Plants expressing Nkd1SRDX 24h after induction of transgene expression (underlined, italics) form a unique group. **b.** Expression pattern of R genes 24h post induction of transgene expression in *A. thaliana* seedlings (Euclidian clustering). Genes depicted are differentially expressed in at least one of the lines compared to the mCherry expressing plants (control). Bold: significantly upregulated in Nkd1. Underlined: significantly upregulated in Nkd1SRDX. Data is the average of 3 independent experiments. Differential gene expression was analyzed with DESeq2, log2Fold change >0.6, p <0.01 (Benjamini-Hochberg correction). **c.** NLR/HR gene expression in maize plants expressing the respective viral constructs was monitored by qRT-PCR and normalized to Zm*CDK* 9dpi. Data is mean ± SD, n=3. No significant differences could be found between the different constructs (p<0.01, Tukey's ANOVA). *LSL3*: *LSD* LIKE3. *ADL2*, *ADR* LIKE2, *LDL1*: *ADR* LIKE1, *TLR1*: ZmTIR-NB-LRR1.

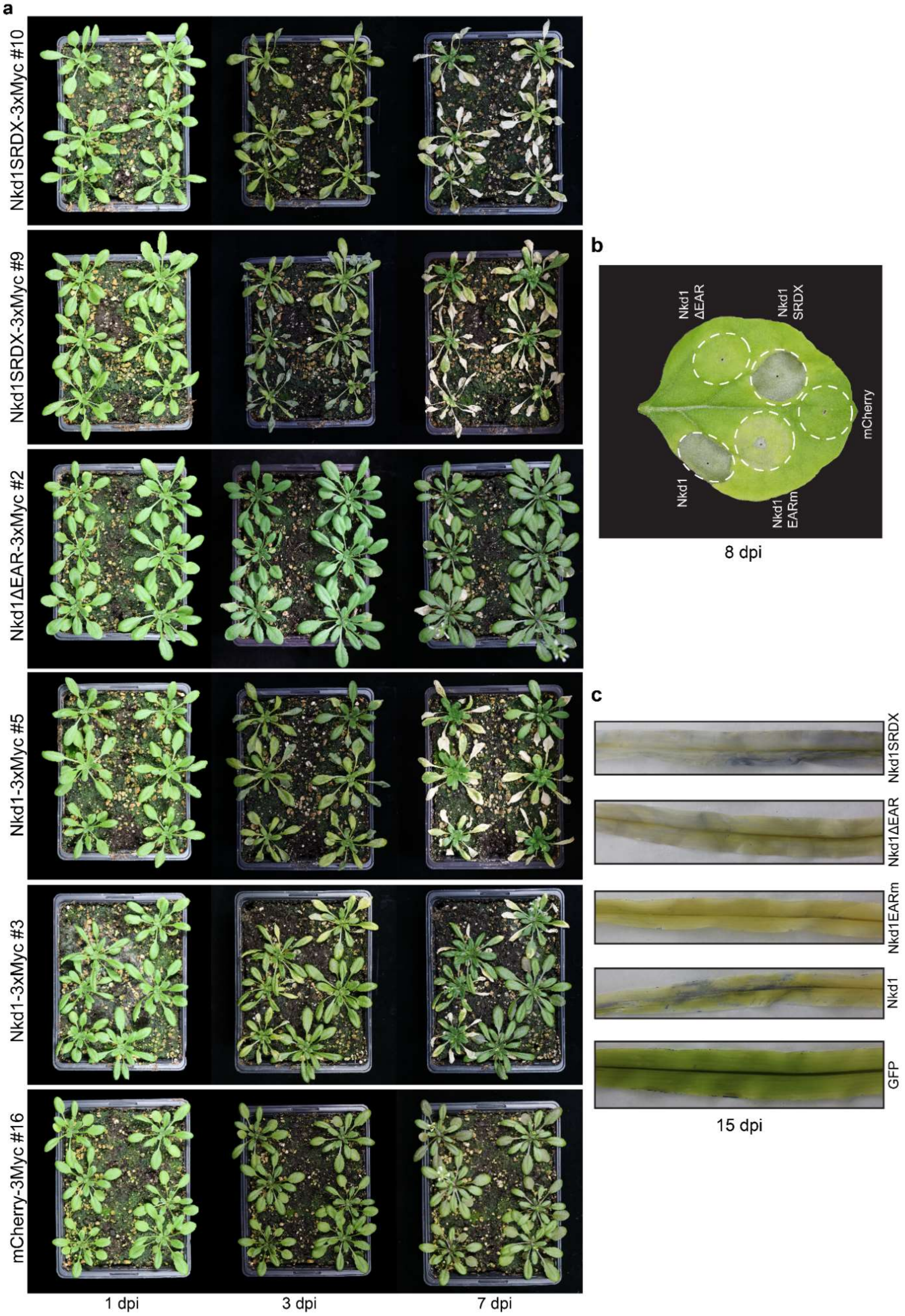

a
Nkd1SRDX-3xMyc #10
Nkd1SRDX-3xMyc #9
Nkd1ΔEAR-3xMyc #2
Nkd1-3xMyc #5
Nkd1-3xMyc #3
mCherry-3Myc #16
1 dpi
3 dpi
7 dpi
b
Nkd1 ΔEAR
Nkd1 SRDX
mCherry
Nkd1
Nkd1 EARm
8 dpi
c
Nkd1SRDX
Nkd1ΔEAR
Nkd1EARm
Nkd1
GFP
15 dpi

**Supplementary Figure S8.**
Cell death phenotype of *A. thaliana* plants expressing different Nkd1 variants. Expression was induced by β-estradiol spray (5 μM) and symptoms were monitored over time. **b.** Cell death phenotype of the different Nkd1 variants in *N. benthamiana*. **c.** Cell death phenotypes of maize plants transiently expressing different Nkd1 variants 15dpi. Plants were stained with Trypan Blue to reveal cell death symptoms.

**Sup. Table1.** Genes differentially expressed in *A. thaliana* seedling expressing different effector constructs 5h post estradiol treatment. Seedlings expressing mCherry were used as the reference. Linked to Fig 4d. n.s.: non-significant

| | | Nkd1ΔEAR | | Nkd1 | | Nkd1SRDX | |
|---|---|---|---|---|---|---|---|
| **Locus** | **Gene name** | **Fold change** | **p-value** | **Fold change** | **p-value** | **Fold change** | **p-value** |
| AT4G16780 | HB-2 | 8.1417426611002 | 4.94378840127362e-07 | n.s. | n.s. | 3.49767859632619 | 0.000979838932320174 |
| AT4G13770 | CYP83A1 | n.s. | n.s. | 2.40010126489914 | 0.000367329544308804 | n.s. | n.s. |
| AT5G53590 | AT5G53590 | 2.18479907595526 | 3.65798210915322e-10 | 1.6390367220084 | 0.00011952699754779 | 0.510635227111713 | 2.59672822989508e-08 |
| AT3G50060 | MYB77 | n.s. | n.s. | n.s. | n.s. | 2.45993149271091 | 0.000234588560792837 |
| AT4G16690 | MES16 | n.s. | n.s. | n.s. | n.s. | 2.45524635581907 | 3.17119790465885e-05 |
| AT2G34650 | PID | n.s. | n.s. | 1.9349785506638 | 0.00288342873534323 | 2.29990452830772 | 1.15227502731234e-05 |
| AT5G13370 | GH3.15 | n.s. | n.s. | n.s. | n.s. | 2.74997740760266 | 3.57452079465051e-10 |
| AT5G66350 | SHI | n.s. | n.s. | n.s. | n.s. | 1.94072000327792 | 0.00138720562820878 |
| AT3G11220 | ELO1 | n.s. | n.s. | n.s. | n.s. | 1.61455286867738 | 0.00339474888855069 |
| AT5G65510 | AIL7 | n.s. | n.s. | n.s. | n.s. | 3.99163406316946 | 4.73663246070452e-06 |
| AT3G62100 | IAA30 | n.s. | n.s. | n.s. | n.s. | 3.59742602659108 | 0.00390402212144196 |
| AT3G44300 | NIT2 | n.s. | n.s. | n.s. | n.s. | 2.96748298119646 | 6.4114198434907e-08 |
| AT3G07390 | AIR12 | n.s. | n.s. | 2.4361512948065 | 0.0012544345157288 | n.s. | n.s. |
| AT5G65980 | PILS7 | n.s. | n.s. | 2.57383343604893 | 0.00438238938001204 | n.s. | n.s. |
| AT1G66600 | ABO3 | n.s. | n.s. | 28.7351724466836 | 7.49035713885383e-20 | 62.2553521623344 | 1.61082092289956e-30 |
| AT1G47510 | 5PTASE11 | n.s. | n.s. | n.s. | n.s. | 6.26690222662871 | 4.9119313312034e-05 |
| AT3G03660 | WOX11 | n.s. | n.s. | 4.70402267060706 | 2.39789571410859e-08 | 3.84889038851293 | 6.72703932874631e-07 |
| AT4G36110 | SAUR9 | n.s. | n.s. | 2.55412574282478 | 8.531937775456e-05 | 2.47816099995627 | 4.28343780005068e-05 |

| | | | | | | | |
|---|---|---|---|---|---|---|---|
| AT4G 35020 | RAC3 | n.s. | n.s. | n.s. | n.s. | 1.5916548 8722205 | 0.001649810 96005381 |
| AT4G 23570 | SGT1 A | n.s. | n.s. | 1.6173865 695906 | 0.002201282 1512757 | 4.2345671 5435878 | 5.606377592 85439e-30 |
| AT2G 26740 | SEH | n.s. | n.s. | n.s. | n.s. | 2.1614213 8082538 | 4.502628786 70925e-05 |
| AT1G 51760 | IAR3 | n.s. | n.s. | 2.0183537 0155613 | 5.962627339 9709e-05 | 2.0695859 7418221 | 6.400543838 41665e-06 |
| AT5G 20810 | SAUR 70 | n.s. | n.s. | n.s. | n.s. | 3.1759832 788823 | 0.001474195 94330116 S |
| AT1G 54990 | AXR4 | n.s. | n.s. | n.s. | n.s. | 2 1.5208963 4694864 | 0.000114102 87348959 |
| AT1G 05680 | AT1G 05680 | n.s. | n.s. | n.s. | n.s. | 4.7410663 4591713 | 0.007754749 77641593 |
| AT1G 31880 | BRX | n.s. | n.s. | n.s. | n.s. | 1.9028735 5001667 | 0.000209607 52582134 |
| AT5G 05730 | ASA1 | n.s. | n.s. | 2.3527455 9378905 | 0.000711206 99693711 | n.s. | n.s. |
| AT3G 28415 | ABCB 22 | n.s. | n.s. | 127.30374 407261 | 1.218345632 85627e-05 | 60.239116 5268745 | 0.000129193 35002548 |
| AT5G 01490 | CAX4 | n.s. | n.s. | 4.5089500 2231445 | 3.528987925 89491e-13 | 2.2049995 1937086 | 0.000275729 533434721 |
| AT4G 02520 | GSTF2 | n.s. | n.s. | 2.7386822 8211328 | 0.008424439 94468253 | 3.6770084 9916848 | 4.722973394 95811e-05 |
| AT4G 37900 | AT4G 37900 | n.s. | n.s. | 2.3588235 2005472 | 7.219989836 17534e-06 | 2.4215511 3903952 | 7.713479241 00064e-07 |
| AT4G 37060 | PLP5 | n.s. | n.s. | 24.395339 1872571 | 3.204094140 07251e-25 | 8.1321062 1169345 | 3.330771337 55821e-11 |
| AT2G 20610 | SUR1 | n.s. | n.s. | 2.0103929 4397998 | 0.000139257 88152063 | n.s. | n.s. |
| AT2G 04160 | AIR3 | n.s. | n.s. | 1.7684144 157309 | 0.001918400 89947976 | n.s. | n.s. |
| AT1G 15580 | IAA5 | n.s. | n.s. | n.s. | n.s. | 14.999449 7190704 | 1.759801712 16352e-08 |
| AT2G 01200 | IAA32 | n.s. | n.s. | n.s. | n.s. | 4.5319178 8718405 | 0.000289896 51266228 |
| AT3G 02875 | ILR1 | n.s. | n.s. | 2.6109441 2344617 | 6.031788614 81939e-07 | 2.6747795 3343346 | 6.640801922 32102e-08 |
| AT1G 73590 | PIN1 | n.s. | n.s. | n.s. | n.s. | 1.9772477 0271379 | 0.001988038 69421421 |
| AT4G 32810 | CCD8 | n.s. | n.s. | n.s. | n.s. | 2.9951999 247451 0. | 0010352922 9815689 ca |
| AT5G 14960 | DEL2 | n.s. | n.s. | n.s. | n.s. | 2.3462900 6423596 | 1.753978404 05105e-05 |

| | | | | | | | |
|---|---|---|---|---|---|---|---|
| AT2G 14960 | GH3.1 | n.s. | n.s. | 3.6583289 7938731 | 0.003619100 10753668 | 9.2620172 0403233 | 1.334068659 49075e-09 |
| AT5G 60450 | ARF4 | n.s. | n.s. | n.s. | n.s. | 1.7938750 9612482 | 3.101280619 13727e-0 |
| AT2G 42430 | LBD16 | n.s. | n.s. | n.s. | n.s. | 4.3343854 5889811 | 2.915028074 64338e-12 |
| AT4G 37390 | BRU6 | n.s. | n.s. | 2.2766580 4991057 | 0.001005846 02747106 | 14.957632 4106932 | 4.205145902 14055e-40 |
| AT2G 23170 | GH3.3 | n.s. | n.s. | n.s. | n.s. | 5.0214430 056626 | 1.529233799 17154e-12 |
| AT5G 56030 | HSP81 -2 | n.s. | n.s. | n.s. | n.s. | 2.0426708 0248998 | 4.165872057 45857e-07 |
| AT5G 56010 | HSP81 -3 | n.s. | n.s. | n.s. | n.s. | 1.5590311 6448606 | 0.001331517 22342473 |
| AT3G 52400 | SYP12 2 | n.s. | n.s. | n.s. | n.s. | 2.1070924 457466 | 0.006029833 92450914 |
| AT2G 27690 | CYP94 C1 | n.s. | n.s. | 1.9668166 216252 | 0.000578878 93869285 | 9.0099176 9354327 | 3.127301146 76274e-45 |
| AT1G 30135 | JAZ8 | n.s. | n.s. | 15.548375 776975 | 2.316514248 91925e-05 | 23.621569 7358878 | 9.939527941 98607e-08 |
| AT3G 45140 | LOX2 | n.s. | n.s. | n.s. | n.s. | 3.5890554 9253871 | 5.763852346 23149e-06 |
| AT1G 70700 | TIFY7 | 2.2261451 3277271 | 2.62292960 545789e-07 | 3.1860104 8206486 | 1.760661860 37306e-18 | 3.4311846 6936592 | 2.188988234 00317e-21 |
| AT3G 12500 | HCHIB | 5.4122340 8693088 | 4.23767278 547731e-15 | 4.9979724 0683711 | 6.487327396 88652e-15 | 2.6874684 7122318 | 2.817390690 31324e-06 |
| AT4G 16760 | ACX1 | n.s. | n.s. | 1.5913745 4753851 | 3.196904828 72185e-07 | 1.6920582 5936081 | 7.755688082 86102e-10 |
| AT5G 06870 | PGIP2 | n.s. | n.s. | n.s. | n.s. | 2.7003440 469712 | 3.703401725 89034e-18 |
| AT5G 20900 | JAZ12 | n.s. | n.s. | n.s. | n.s. | 1.5287426 8040871 | 0.001644780 60673348 |
| AT2G 24850 | TAT3 | n.s. | n.s. | n.s. | n.s. | 2.0075815 580033 | 8.937840731 07446e-20 |
| AT5G 26170 | WRKY 50 | n.s. | n.s. | n.s. | n.s. | 23.193516 6909221 | 1.459043981 29141e-11 |
| AT5G 64810 | WRKY 51 | n.s. | n.s. | 14.796914 337714 | 1.045298282 78133e-10 | 21.195999 7950388 | 2.420552217 84457e-14 |
| AT2G 34600 | JAZ7 | n.s. | n.s. | 3.2346383 3689455 | 0.007769081 59488448 | 3.0896881 699082 | 0.003494397 71760999 |
| AT5G 13220 | JAZ10 | n.s. | n.s. | 8.2240176 0852207 | 1.941768651 35545e-53 | 8.4057811 8744586 | 7.286550872 63351e-55 |
| AT1G 74950 | TIFY1 0B | n.s. | n.s. | 3.1303547 9541919 | 2.252709318 56733e-45 | 2.9762187 6143948 | 2.055727219 79637e-41 |

| | | | | | | | |
|---|---|---|---|---|---|---|---|
| AT1G32640 | MYC2 | n.s. | n.s. | 2.0182050511116 | 6.47918183454665e-08 | 2.10401537501309 | 1.76266124259169e-09 |
| AT1G19180 | JAZ1 | n.s. | n.s. | 2.76517403492804 | 1.1003002946804e-06 | 1.81688081414846 | 0.00480759411246093 |
| AT2G06050 | OPR3 | n.s. | n.s. | 2.26998308040087 | 2.00959097747345e-11 | 1.66573501322098 | 4.61148772094076e-05 |
| AT1G55020 | LOX1 | n.s. | n.s. | 2.18432843653976 | 4.92907984002056e-05 | n.s. | n.s. |
| AT5G42650 | AOS | n.s. | n.s. | 2.03748492972354 | 0.000787512468747723 | n.s. | n.s. |
| AT3G55970 | JRG21 | n.s. | n.s. | 9.85814593325012 | 9.47475097856404e-15 | n.s. | n.s. |
| AT5G24770 | VSP2 | n.s. | n.s. | 3.17018676535551 | 0.000516920018761546 | n.s. | n.s. |
| AT3G17860 | JAZ3 | n.s. | n.s. | 2.25228032519773 | 1.20360295792371e-05 | n.s. | n.s. |
| AT5G18860 | NSH3 | n.s. | n.s. | 1.5250190812911 | 0.00158919873172406 | 0.640084654606449 | 0.00019119730099225 |
| AT3G30775 | ERD5 | n.s. | n.s. | n.s. | n.s. | 3.3334572554761 | 7.98539063911977e-07 |
| AT2G39030 | NATA1 | n.s. | n.s. | 93.328347638866 | 0.00238059009060659 | 75.5512788199907 | 0.00118175421662614 Ac |
| AT4G11650 | OSM34 | n.s. | n.s. | 7.69623803048736 | 1.97822552859635e-19 | 4.88195302008801 | 2.79713442123534e-12 |
| AT3G19580 | ZF2 | n.s. | n.s. | 3.94643571117453 | 2.70098169464734e-14 | 2.23627705527 | 1.5192318909492e-05 |
| AT1G52890 | NAC019 | n.s. | n.s. | 4.77147785197416 | 0.00132319142672779 | 6.69704919635566 | 6.27898438034249e-06 |
| AT1G15520 | ABCG40 | n.s. | n.s. | 10.3154053112058 | 3.19941986588103e-05 | 24.9770494453903 | 1.87746413154377e-10 |
| AT3G51810 | EM1 | n.s. | n.s. | 41.5856598840345 | 0.00190966088448411 | n.s. | n.s. |
| AT2G40180 | PP2C5 | n.s. | n.s. | n.s. | n.s. | 11.8788997066928 | 6.66263452013914e-0 |
| AT2G37770 | ChlAKR | n.s. | n.s. | n.s. | n.s. | 11.0962782594279 | 9.01607251292409e-23 |
| AT2G36270 | ABI5 | n.s. | n.s. | n.s. | n.s. | 4.28467173070934 | 5.37020879417963e-07 B |
| AT2G41010 | CAMBP25 | n.s. | n.s. | 3.52598672022503 | 3.42614326138463e-06 | 3.96600195762145 | 5.41113164636741e-08 |
| AT1G01720 | ATAF1 | n.s. | n.s. | 2.30925307611389 | 0.000319814691678936 | 2.96216470469458 | 1.34126164811e-07 |
| AT2G38905 | AT2G38905 | n.s. | n.s. | n.s. | n.s. | 11.1842365347703 | 9.58746955369077e-06 |
| AT3G24650 | ABI3 | n.s. | n.s. | n.s. | n.s. | 6.5921351476252 | 3.86010734881655e-05 |

| | | | | | | | |
|---|---|---|---|---|---|---|---|
| AT4G 28520 | CRU3 | n.s. | n.s. | n.s. | n.s. | 61.645338 4235471 | 0.001283923 41205128 |
| AT5G 44120 | CRA1 | n.s. | n.s. | n.s. | n.s. | 31.959763 1887576 | 0.000889305 283638038 |
| AT1G 03880 | CRU2 | n.s. | n.s. | n.s. | n.s. | 126.83439 8616583 | 0.000432905 80282777 |
| AT1G 18100 | E12A1 1 | n.s. | n.s. | 4.1361246 0988632 | 7.094379452 27984e-11 | 15.152426 0298121 | 7.842222685 59156e-41 |

**Sup. Table 2.** Genes differentially expressed in *A. thaliana* seedling expressing different effector constructs 24h post estradiol treatment. Seedling expressing mCherry were used as the reference. Linked to Fig 6d. n.s: non-significant

| | | Nkd1ΔEAR | | Nkd1 | | Nkd1SRDX | |
|---|---|---|---|---|---|---|---|
| **Locus** | **Gene name** | **Fold change** | **p-value** | **Fold change** | **p-value** | **Fold change** | **p-value** |
| AT1G70710 | GH9B1 | n.s. | n.s. | n.s. | n.s. | 1.75802341036277 | 0.00925283707252297 |
| AT2G14580 | PRB1 | n.s. | n.s. | n.s. | n.s. | 3.31623068270488 | 0.00295415863697124 |
| AT5G25980 | TGG2 | n.s. | n.s. | n.s. | n.s. | 2.23339689341286 | 1.1810542908005e-07 |
| AT2G19970 | AT2G19970 | n.s. | n.s. | n.s. | n.s. | 3.95432212599826 | 8.26456092590891e-08 |
| AT3G45300 | IVD | n.s. | n.s. | n.s. | n.s. | 2.19026043496475 | 0.000522575344728425 |
| AT2G02120 | PDF2.1 | n.s. | n.s. | n.s. | n.s. | 8.53140552452 | 2.35999881012556e-06 |
| AT5G43570 | AT5G43570 | n.s. | n.s. | n.s. | n.s. | 4.79799371151703 | 0.000314962023587233 |
| AT2G19990 | PR-1-LIKE | n.s. | n.s. | n.s. | n.s. | 2.37512741897822 | 0.00176641720257293 |
| AT3G54420 | EP3 | n.s. | n.s. | n.s. | n.s. | 4.16241989571996 | 1.75502443858664e-18 |
| AT2G02100 | LCR69 | n.s. | n.s. | n.s. | n.s. | 1.96227092477496 | 0.00016451733998571 |
| AT2G15010 | AT2G15010 | n.s. | n.s. | n.s. | n.s. | 355.790801327728 | 1.57448216387526e-07 |
| AT3G57240 | BG3 | n.s. | n.s. | 6.61154978500811 | 0.00023372432147690 | 643.032723782415 | 7.42785705630819e-51 |
| AT2G14610 | PR1 | n.s. | n.s. | n.s. | n.s. | 1056.61922741197 | 1.85363080863306e-13 |
| AT2G38900 | AT2G38900 | n.s. | n.s. | n.s. | n.s. | 151.692355464302 | 2.6888918383293e-05 |
| AT4G24260 | GH9A3 | n.s. | n.s. | 3.40016349608637 | 0.0002161369555830 | 6.05017434243293 | 4.18786164618454e-10 |
| AT5G59320 | LTP3 | n.s. | n.s. | n.s. | n.s. | 2.93742945068868 | 0.00302188955108147 |
| AT1G52030 | MBP2 | n.s. | n.s. | 3.94040143216126 | 0.0002474701382314 | 5.64539269383546 | 1.57712529469985e-07 |
| AT1G75040 | PR5 | n.s. | n.s. | 3.82535774621893 | 0.00083944900316847 | 10.2452337615628 | 1.32352318290645e-11 |
| AT4G07820 | AT4G07820 | n.s. | n.s. | 5.24630999941535 | 9.92504349146828e-14 | 32.3311493480779 | 6.20022139370454e-63 |

| | | | | | | | |
|---|---|---|---|---|---|---|---|
| AT4G01700 | AT4G01700 | 2.4697099775008 | 0.0020321245841066 | 11.4369144870395 | 1.01176115374289e-26 | 62.1376567839578 | 1.29767091865007e-78 |
| AT3G52430 | PAD4 | n.s. | n.s. | n.s. | n.s. | 29.3430989304656 | 7.59764388142912e-51 |
| AT5G24620 | AT5G24620 | n.s. | n.s. | n.s. | n.s. | 1.63683746390844 | 0.00790871124426115 |
| AT1G78780 | AT1G78780 | n.s. | n.s. | n.s. | n.s. | 8.02198206947604 | 9.80115776961581e-25 |
| AT3G47540 | AT3G47540 | n.s. | n.s. | 6.04882116417854 | 1.70960683102522e-14 | 46.8036698498645 | 9.42074304927876e-68 |
| AT3G57260 | BGL2 | n.s. | n.s. | n.s. | n.s. | 79.0234858250542 | 1.35578750330484e-06 |
| AT1G62540 | FMO GS-OX2 | n.s. | n.s. | n.s. | n.s. | 3.8331308373683 | 6.85399912404822e-07 |
| AT3G47960 | GTR1 | n.s. | n.s. | n.s. | n.s. | 2.1194930146908 | 1.25725567140593e-07 |
| AT2G43570 | CHI | n.s. | n.s. | n.s. | n.s. | 4.44798515829311 | 8.63282407421244e-08 |
| AT4G19810 | ChiC | n.s. | n.s. | 1.97161368544448 | 0.0057629446817065 | 6.30492077758618 | 1.21293543394828e-20 |
| AT1G02360 | AT1G02360 | n.s. | n.s. | n.s. | n.s. | 6.89319358013723 | 1.48087129338798e-18 |
| AT3G04720 | PR4 | n.s. | n.s. | n.s. | n.s. | 3.73313870763541 | 4.85576807463897e-12 |
| AT4G33720 | AT4G33720 | n.s. | n.s. | 70.2586456544199 | 0.0016451719299178 | 3.0100685104847 | 0.000522575344728425 |
| AT1G50050 | AT1G50050 | n.s. | n.s. | 52.9405298778398 | 5.49411637262414e-09 | n.s. | n.s. |
| AT2G43590 | AT2G43590 | n.s. | n.s. | 2.76399387394929 | 6.01281867096194e-10 | n.s. | n.s. |
| AT4G19750 | AT4G19750 | n.s. | n.s. | 146.930577684256 | 1.75010938479313e-06 | 45.6583524320388 | 0.000121360920613606 |
| AT4G19760 | AT4G19760 | n.s. | n.s. | 277.280769962848 | 0.000105967254311474 | 48.3540730473376 | 0.00485563854633187 |
| AT4G19720 | AT4G19720 | n.s. | n.s. | 6.04741437018495 | 8.65177207919521e-13 | 2.26619768132522 | 0.00161509298137455 |
| AT3G57270 | PAD3 | n.s. | n.s. | 68.9544456864653 | 0.0033178027247168 | n.s. | n.s. |
| AT3G26830 | BG1 | n.s. | n.s. | 13.3331548299754 | 2.83301075310879e-0 | 6.39396074234553 | 0.000435896135686253 |

**Sup. Table3.** Genes differentially expressed in *A. thaliana* seedling expressing different effector constructs 24h post estradiol treatment. Seedling expressing mCherry were used as the reference. Linked to Fig S7b. n.s: non-significant

| | | **Nkd1ΔEAR** | | **Nkd1** | | **Nkd1SRDX** | |
|---|---|---|---|---|---|---|---|
| **Locus** | **Gene name** | **Fold change** | **p-value** | **Fold change** | **p-value** | **Fold change** | **p-value** |
| AT5G 41550 | AT5G 41550 | n.s. | n.s. | n.s. | n.s. | 3.2605187 7433731 | 0.00039086 278073318 |
| AT1G 72850 | AT1G 72850 | n.s. | n.s. | n.s. | n.s. | 3.8850796 3989699 | 2.41889850 164119e-07 |
| AT1G 47370 | AT1G 47370 | n.s. | n.s. | n.s. | n.s. | 7.8185129 2387935 | 5.88817873 970206e-10 |
| AT3G 44400 | AT3G 44400 | n.s. | n.s. | 2.1358282 5372085 | 2.98776304 211345e-05 | 6.9313193 5514667 | 7.94302167 271874e-35 |
| AT5G 46520 | AT5G 46520 | n.s. | n.s. | 3.7526440 9205851 | 1.79889367 306854e-07 | 14.916801 5660019 | 1.90200876 332465e-33 |
| AT5G 46260 | AT5G 46260 | n.s. | n.s. | 3.8118778 6507912 | 6.37342086 172382e-09 | 15.832133 1647862 | 1.17945189 902704e-4 |
| AT5G 45090 | PP2-A7 | n.s. | n.s. | 9.7486349 955783 | 0.00044930 638875625 | 50.885420 1686218 | 9.03810442 259590e-13 |
| AT5G 41740 | AT5G 41740 | n.s. | n.s. | 2.3658291 3989538 | 0.00232036 911427453 | 9.5037737 6386663 | 3.48614894 859751e-22 |
| AT5G 45000 | AT5G 45000 | n.s. | n.s. | 3.1151351 430275 | 8.20833088 073197e-05 | 14.199706 9333624 | 7.61334632 718956e-27 |
| AT3G 44670 | AT3G 44670 | n.s. | n.s. | n.s. | n.s. | 2.9651315 448973 | 1.34725307 728249e-14 |
| AT5G 46490 | AT5G 46490 | n.s. | n.s. | n.s. | n.s. | 5.0227070 7508222 | 7.07638712 153579e-15 |
| AT5G 45240 | AT5G 45240 | n.s. | n.s. | 4.0993795 4526403 | 0.00557815 30804782 | 56.209148 3706826 | 3.90153223 194397e-25 |
| AT1G 56520 | AT1G 56520 | n.s. | n.s. | n.s. | n.s. | 4.7455391 1724537 | 1.61787336 924524e-33 |
| AT5G 18350 | AT5G 18350 | n.s. | n.s. | n.s. | n.s. | 7.8971335 8890118 | 1.39515184 473562e-09 |
| AT4G 16860 | AT4G 16860 | n.s. | n.s. | n.s. | n.s. | 7.8971335 8890118 | 1.39515184 473562e-09 |
| AT1G 57630 | AT1G 57630 | n.s. | n.s. | n.s. | n.s. | 87.870547 1273566 | 3.85831499 517556e-12 |
| AT1G 64070 | RLM1 | n.s. | n.s. | n.s. | n.s. | 106.20391 719982 | 1.11026729 123736e-05 |
| AT5G 38344 | AT5G 38344 | n.s. | n.s. | n.s. | n.s. | 14.414511 5977973 | 9.90623704 454447e-06 |
| AT1G 63350 | AT1G 63350 | n.s. | n.s. | n.s. | n.s. | 4.8519481 2143731 | 1.08251004 69745e-06 |

| | | | | | | | |
|---|---|---|---|---|---|---|---|
| AT1G 51270 | AT1G 51270 | n.s. | n.s. | n.s. | n.s. | 12.594922 4443167 | 1.54913718 795238e-17 |
| AT1G 66090 | AT1G 66090 | n.s. | n.s. | 2.4012629 7759565 | 0.00238073 321455307 | 14.140854 0655303 | 1.78115316 394951e-29 |
| AT3G 25510 | AT3G 25510 | n.s. | n.s. | n.s. | n.s. | 13.893500 667977 | 9.91840424 851349e-21 |
| AT3G 04220 | AT3G 04220 | n.s. | n.s. | 8.1707794 4496172 | 1.75980354 029921e-09 | 58.548752 9065786 | 3.62424050 670793e-37 |
| AT2G 32140 | AT2G 32140 | 3.4699317 8383742 | 0.00439815 697318625 | 14.829561 8745094 | 9.38084720 162036e-17 | 102.63212 5478673 | 1.42934070 438609e-51 |
| AT4G 16890 | SNC1 | n.s. | n.s. | n.s. | n.s. | 3.2327941 3469657 | 7.95078639 232061e-28 |
| AT1G 17600 | AT1G 17600 | n.s. | n.s. | 2.8196551 1801335 | 0.00501089 020186444 | 14.941346 285846 | 4.95678222 997693e-20 |
| AT2G 20142 | AT2G 20142 | n.s. | n.s. | 3.7971021 5746312 | 0.00054178 089903433 | 23.534058 2815137 | 4.19800313 56908e-22 |
| AT2G 17050 | AT2G 17050 | n.s. | n.s. | n.s. | n.s. | 21.360665 9802503 | 3.84788822 332427e-1 |
| AT5G 38350 | AT5G 38350 | n.s. | n.s. | 2.3571692 2603986 | 0.00057841 084332309 | 11.437550 8409312 | 8.47205394 575417e-33 |
| AT5G 04720 | ADR1-L2 | n.s. | n.s. | n.s. | n.s. | 5.0056851 6546168 | 3.46896225 967467e-41 |
| AT1G 51480 | AT1G 51480 | n.s. | n.s. | n.s. | n.s. | 5.1663357 4798465 | 0.00656005 702241929 |
| AT1G 33560 | AT1G 33560 | n.s. | n.s. | n.s. | n.s. | 3.5670865 1841563 | 8.36195152 254505e-18 |
| AT5G 58120 | AT5G 58120 | n.s. | n.s. | n.s. | n.s. | 3.4204751 4505382 | 9.59579121 047872e-13 |
| AT1G 15890 | AT1G 15890 | n.s. | n.s. | n.s. | n.s. | 4.1288587 4473212 | 3.48865843 627404e-11 |
| AT4G 19520 | AT4G 19520 | n.s. | n.s. | 1.9191491 3963059 | 6.18963857 583529e-05 | 4.7576318 1453348 | 1.25687170 034377e-28 |
| AT2G 16870 | AT2G 16870 | n.s. | n.s. | n.s. | n.s. | 4.2776390 7714657 | 1.21702785 98247e-13 |
| AT1G 59780 | AT1G 59780 | n.s. | n.s. | 4.7556595 7317612 | 4.08650458 522045e-06 | 13.895793 4210321 | 2.71897204 963125e-18 |
| AT1G 17610 | AT1G 17610 | n.s. | n.s. | n.s. | n.s. | 5.2510833 3737654 | 6.14452635 377523e-07 |
| AT5G 38340 | AT5G 38340 | n.s. | n.s. | 2.6035415 6370715 | 9.69829231 34646e-07 | 6.8723924 5252266 | 1.44303038 088626e-28 |
| AT1G 72900 | AT1G 72900 | n.s. | n.s. | 7.4001390 824336 | 4.80142681 31582e-14 | 23.918396 968909 | 1.87124556 096437e-36 |
| AT1G 72930 | TIR | n.s. | n.s. | 4.4965832 9082733 | 4.12424748 372085e-12 | 13.724515 3368611 | 7.60626463 307359e-38 |
| AT1G 17615 | AT1G 17615 | n.s. | n.s. | 77.002799 6486259 | 0.00082522 91151027 | 291.85235 6878207 | 5.12861632 317149e-0 |

| | | | | | | | |
|---|---|---|---|---|---|---|---|
| AT4G 33300 | ADR1-L1 | n.s. | n.s. | 2.0244804 7973114 | 8.51628993 57568e-07 | 5.4061818 4647729 | 1.67450569 784458e-39 |
| AT5G 41750 | AT5G 41750 | n.s. | n.s. | 5.8591202 8175217 | 1.47777394 255875e-05 | 20.203287 426122 | 1.32846038 696825e-16 |
| AT1G 63750 | AT1G 63750 | n.s. | n.s. | n.s. | n.s. | 3.1066128 1912135 | 1.47751053 232036e-14 |
| AT3G 44630 | AT3G 44630 | n.s. | n.s. | n.s. | n.s. | 3.0409920 752845 | 2.12018634 153734e-18 |
| AT4G 14370 | AT4G 14370 | n.s. | n.s. | 3.0460001 3514112 | 0.00035599 595762225 | 10.324301 1618105 | 2.16312762 778199e-18 |
| AT1G 31540 | AT1G 31540 | n.s. | n.s. | n.s. | n.s. | 3.1707800 2217717 | 4.97980681 780667e-13 |
| AT4G 16960 | AT4G 16960 | n.s. | n.s. | 2.3396110 9881959 | 4.57286891 179307e-05 | 7.5596372 4472119 | 1.85942485 891707e-29 |
| AT3G 14470 | AT3G 14470 | n.s. | n.s. | n.s. | n.s. | 2.9736236 061397 | 8.11553445 509481e-14 |
| AT4G 11170 | AT4G 11170 | n.s. | n.s. | 17.627107 6371782 | 1.76772318 383355e-09 | 84.013155 3740526 | 1.70559293 667869e-23 |
| AT3G 04210 | AT3G 04210 | n.s. | n.s. | 3.6694133 3626178 | 3.24337022 384885e-13 | 13.947295 8417417 | 5.07092788 243501e-56 |
| AT4G 19920 | AT4G 19920 | n.s. | n.s. | 12.230408 0090915 | 1.39288726 890778e-14 | 53.720933 081118 | 1.99972148 915496e-38 |
| AT4G 26090 | RPS2 | n.s. | n.s. | 2.3945522 5306658 | 0.00030474 087894697 | 7.4393546 7651049 | 2.15436823 935914e-22 |
| AT5G 66630 | DAR5 | n.s. | n.s. | 3.2034693 2523743 | 5.69754149 976833e-06 | 11.059050 2732341 | 2.01801186 652403e-26 |
| AT5G 46470 | RPS6 | n.s. | n.s. | n.s. | n.s. | 3.3725815 595556 | 1.21520712 314074e-18 |
| AT3G 50950 | ZAR1 | n.s. | n.s. | n.s. | n.s. | 3.0685747 9606036 | 6.32440013 380549e-10 |
| AT1G 12210 | RFL1 | n.s. | n.s. | n.s. | n.s. | 4.2234945 4379583 | 2.50250819 228172e-10 |
| AT1G 72910 | AT1G 72910 | n.s. | n.s. | 11.301817 3038356 | 2.55342121 820112e-10 | 31.891164 9041892 | 4.67628890 336782e-22 |
| AT1G 50180 | AT1G 50180 | n.s. | n.s. | n.s. | n.s. | 3.6828583 2405099 | 0.00078292 187262625 |
| AT4G 19925 | AT4G 19925 | n.s. | n.s. | 20.970909 2555661 | 1.92278168 586494e-08 | 49.724395 8902974 | 7.49788376 724712e-15 |
| AT1G 72940 | AT1G 72940 | n.s. | n.s. | 2.2084218 7514585 | 0.00013312 968703456 | 4.2481777 0740175 | 1.08775203 603653e-15 |
| AT1G 52900 | AT1G 52900 | n.s. | n.s. | n.s. | n.s. | 53.580928 3492856 | 0.00230329 200178562 |
| AT5G 18370 | AT5G 18370 | n.s. | n.s. | 1.6949506 921745 | 0.00109548 216743356 | 2.5201412 6822722 | 1.33219981 867876e-11 |
| AT1G 72950 | AT1G 72950 | n.s. | n.s. | n.s. | n.s. | 4.4742064 5608356 | 4.43515260 561374e-0 |

| | | | | | | | |
|---|---|---|---|---|---|---|---|
| AT5G 45080 | PP2-A6 | 6.91778256876665 | 9.61994444454162e-05 | 15.7716509899015 | 9.64358880492599e-11 | 45.8493063806659 | 8.29962855839294e-22 |
| AT3G 51560 | AT3G 51560 | n.s. | n.s. | n.s. | n.s. | 59.588196769112 | 5.64315029709698e-05 |
| AT1G 72890 | AT1G 72890 | n.s. | n.s. | n.s. | n.s. | 3.54802329005392 | 2.60243982391832e-10 |
| AT4G 36150 | AT4G 36150 | n.s. | n.s. | 2.29815835109104 | 2.66198557456531e-08 | 6.30808161461921 | 6.25500440224169e-43 |
| AT1G 56540 | AT1G 56540 | n.s. | n.s. | n.s. | n.s. | 4.73706952226184 | 3.27618503937612e-09 |
| AT4G 23515 | AT4G 23515 | n.s. | n.s. | 13.4802520517853 | 4.68720184031973e-11 | 7.22329648306319 | 4.43616694877439e-07 |
| AT4G 27220 | AT4G 27220 | n.s. | n.s. | 12.316775055931 | 0.00593714774478473 | n.s. | n.s. |
| AT5G 45490 | AT5G 45490 | n.s. | n.s. | 1.71845110595735 | 2.34764228 81267e-05 | 1.81918848611606 | 3.06428080877246e-0 |
| AT5G 47250 | AT5G 47250 | n.s. | n.s. | 2.42148804136523 | 0.00016965903323041 | 2.63757076150859 | 4.97769487992395e-06 |
| AT1G 72920 | AT1G 72920 | n.s. | n.s. | 5.48207578771955 | 3.90106138590921e-06 | 6.80830466117086 | 1.58233276587661e-08 |
| AT5G 45070 | PP2-A8 | n.s. | n.s. | 3.00970056964034 | 0.00020614390814045 | 3.64925517859363 | 1.04662680331417e-06 |
| AT5G 44920 | AT5G 44920 | n.s. | n.s. | 3.10218817292371 | 4.20363766919568e-08 | 4.11600093577081 | 1.87696233853363e-13 |
| AT5G 66910 | AT5G 66910 | n.s. | n.s. | 1.86686861018111 | 0.0005032797855819 | 2.2202281645008 | 4.38206194137978e-07 |
| AT4G 16920 | AT4G 16920 | n.s. | n.s. | 3.72039584753558 | 4.27336527849264e-06 | 5.57878717729207 | 3.56765621331524e-11 |
| AT4G 11340 | AT4G 11340 | n.s. | n.s. | 87.8971470135997 | 0.0001937287394422 | 130.126469208441 | 6.23957569430284e-06 |
| AT5G 11250 | AT5G 11250 | n.s. | n.s. | 1.75347402476911 | 0.0006043538553081 | 2.13371244056004 | 1.22131919663101e-07 |
